# Shoot-to-root mobile CEPD proteins regulate TGA transcription factors to allow nitrate root-to-shoot transport in *Arabidopsis thaliana*

**DOI:** 10.1101/2023.10.18.562952

**Authors:** Anja Maren Pelizaeus, Corinna Thurow, Lisa Oskam, Ben Moritz Hoßbach, Jelena Budimir, Ronald Pierik, Christiane Gatz

**Affiliations:** Albrecht-von-Haller-Institut für Pflanzenwissenschaften, Georg-August-Universität Göttingen, Julia-Lermontowa-Weg 3, D-37077 Göttingen, Germany; Plant-Environment Signaling, Dept. Biology, Utrecht University, Padualaan 8, 3584 CH, Utrecht, The Netherlands

## Abstract

In *Arabidopsis thaliana*, nitrogen (N) starvation leads to increased synthesis of CEPD (C-TERMINALLY ENCODED PEPTIDE DOWNSTREAM) proteins in the shoot. CEPDs travel to the roots, where they activate expression of genes required for high affinity nitrate transport. CEPDs belong to a plant-specific class of glutaredoxin-like proteins that interact with TGACG-binding transcription factors (TGAs). Here we identified the redundant clade-I TGAs TGA1 and TGA4 as the link between CEPDs and target promoters. In the absence of CEPDs, TGA1/4 have a strong negative effect on N starvation-induced gene expression leading to reduced translocation of N from the root to the shoot and to reduced shoot fresh weight. Basal levels of CEPDs were sufficient to completely release TGA1/4-mediated repression of nitrate acquisition. The antagonism between CEPDs and TGA1/4 was also detected in shoots, where CEPDs dampened the activating function of TGA1/4 on hyponasty and defense. CEPDs encode the conserved putative active site motif CxxC/S that was suggested to mediate redox regulation of target proteins. Complementation of the *tga1 tga4* mutant with a TGA1 variant containing amino acid substitutions of all four potentially redox-active cysteines showed that CEPDs do not regulate TGA1/4 by modulating their redox state.

## Introduction

As autotrophic organisms, plants depend on the uptake of inorganic molecules like carbon dioxide, water and mineral nutrients. Nitrogen (N) is an essential macronutrient that is available in the soil as nitrate due to bacterial nitrification of reduced organic N. The efficiency of N uptake and assimilation is well adjusted to N availability (Fredes et al., 2019) and constitutes an important trait of crop plants. At least in the model plant *Arabidopsis thaliana*, one of the underlying regulatory mechanisms for N acquisition under N starvation involves the action of C-TERMINALLY ENCODED PEPTIDE DOWNSTREAM (CEPD) proteins, which belong to the family of plant-specific glutaredoxin-like proteins (Ohkubo et al., 2017; Ota et al., 2020; Ohkubo et al., 2021).

Glutaredoxins are small proteins (12-15 kDa) found in all domains of life. The founding member of this protein family acts as an oxidoreductase that requires the atypical tripeptide glutathione as an acceptor or donor of hydrogen (Laurent et al., 1964). As summarized in several reviews (Lillig et al., 2008; Rouhier et al., 2008; Couturier et al., 2009; Lillig and Berndt, 2013; Couturier et al., 2015), glutaredoxins are characterized by the conserved thioredoxin fold positioning the active site with the catalytic cysteine at the N terminus of αhelix 1. Based on their ability to act as oxidoreductases on small glutathionylated substrates like bis(2-hydroxyethyl)disulfide (HEDS) or glutathionylated cysteine, glutaredoxins have been grouped into two classes (Liedgens et al., 2020). Class I glutaredoxins, which contain variants of the CPYC motif in their active sites, are catalytically active, while class II glutaredoxins, which contain a highly conserved CGFS motif, have only negligible GSH-dependent oxidoreductase activity. Instead, they bind labile [2Fe-2S] clusters for transfer to client proteins (Rodriguez-Manzaneque et al., 2002).

According to sequence alignments, land plants contain a group of glutaredoxin-like proteins, which are called CC-type glutaredoxin-like proteins because of their conserved CCM/LC or CCM/LS motif (Rouhier et al., 2008). The few biochemical studies that are available have unraveled either weak (Couturier et al., 2010) or no oxidoreductase activity in the standard HEDS assay (Xu et al., 2022). Since it is not known whether they exert functions related to known glutaredoxin-related functions, we call them glutaredoxin-like proteins. The family of CC-type glutaredoxin-like proteins has expanded during land plant evolution and is represented by 21 members (called ROXYs) in *Arabidopsis thaliana* (Ziemann et al., 2009). They are located in the cytosol and the nucleus (Li et al., 2009; Ohkubo et al., 2017; Xu et al., 2022) and interact with TGACG-binding transcription factors (TGAs) (Ndamukong et al., 2007; Li et al., 2009; Yang et al., 2021). Epistasis analysis has proven the functional relevance of the interaction of CC-type glutaredoxin-like proteins and TGAs: Arabidopsis ROXY1 represses the negative activity of TGA transcription factor PERIANTHIA (PAN) on petal primordia initiation (Li et al., 2009) and the maize CC-type glutaredoxin MSCA1 and its two orthologues ZmGRX2 and ZmGRX5 regulate the activity of TGA factor FEA4 in order to control meristem size (Yang et al., 2021).

Recently, members of a subclade of ROXYs (ROXY6, ROXY7, ROXY8, ROXY9) have been found to be important for nitrate acquisition on medium with high, moderate and low N supply (Ota et al., 2020). These ROXYs have been called CEPD or CEPD-like (CEPDL), because two of them act DOWNSTREAM (D) of C-TERMINALLY ENCODED PEPTIDE (CEP) (Ohkubo et al., 2017). CEP is an N starvation-induced 15-amino-acid peptide hormone, which acts as a root derived N-demand signal. CEPs are transported to the shoot, where they activate signaling through the leucine rich receptor kinases CEP RECEPTOR (CEPR) 1 and 2 (Tabata et al., 2014) leading to increased expression of *CEPD1* alias *ROXY6* and *CEPD2* alias *ROXY9* (Ohkubo et al., 2017). *CEPDL2* alias *ROXY8* is induced upon sensing the N status of the shoot (Ota et al., 2020). CEPDs travel through the phloem to the roots and initiate the expression of genes involved in nitrate uptake like e.g. the nitrate transporter *NRT2.1* and phosphatase *CEPH* (Ohkubo et al., 2017; Ota et al., 2020; Ohkubo et al., 2021) the gene product of which enhances NRT2.1 transport activity by dephosphorylation (Ohkubo et al., 2021). Importantly, nitrate acquisition is strongly impaired in *cepd* mutants leading to reduced fresh weight even under sufficient N supply (Ota et al., 2020; Ohkubo et al., 2021).

Analysis of publicly available expression data led to the hypothesis that clade-I TGAs play a role in the nitrate response of *Arabidopsis thaliana*, although nitrate uptake is not impaired in the *tga1 tga4* mutant (Alvarez et al., 2014). TGA1/4 was subsequently shown to be part of a nitrate-induced signaling cascade leading to increased root hair density (Canales et al., 2017) and to confer N dose-responsive gene expression (Swift et al., 2020). Here, we tested whether CEPDs regulate TGA1/4 activity in the context of the N starvation response. We show that TGA1/4 mediate transcriptional repression on a set of N transport-related genes, some of them being required for efficient N root to shoot allocation. CEPDs are required to release this repression. We further demonstrate that the shoot-borne functions of TGA1/4 as activators of hyponastic growth (Li et al., 2019) and of defense responses (Sun et al., 2018) are dampened by CEPDs, providing other examples for weak but detectable antagonistic interactions of CEPDs and TGA1/4.

## Results

### CEPDs interfere with the repressive action of TGA1 and TGA4 on shoot biomass accumulation

In order to explore whether CEPDs regulate TGA1/4 activity, we planned to perform epistasis analyses with loss of function *cepd* and *tga1 tga4* plants. However, the *tga1 tga4* alleles are in the Col-0 ecotype (Shearer et al., 2012), while *cepd* alleles are in Noessen (Ota et al., 2020). In order to avoid inhomogeneous genetic backgrounds in the planned hexuple mutant, we first generated a *cepd* mutant in Col-0 by CRISPR/Cas9-based genome editing (Supplemental Fig. 1). Since defects in nitrate acquisition and growth of the Noessen ecotype were most pronounced when all four *CEPD* genes were mutated (Ota et al., 2020), we focussed on the analysis of the *roxy6 roxy7 roxy8 roxy9* quadruple mutant, which we will call *cepd* from here on. When grown on fertilized soil, the fresh weight of *cepd* was reduced by 53% under long day conditions (16L/8D) and by 47% under a 12L/12D cycle (Fig. 1). The homozygous progeny of the cross between *cepd* and *tga1 tga4* accumulated the same amount of fresh weight as Col-0 and *tga1 tga4*. Apparently, TGA1/4 have a strong growth-repressing function that is completely relieved through the action of CEPDs.

**Fig. 1.**
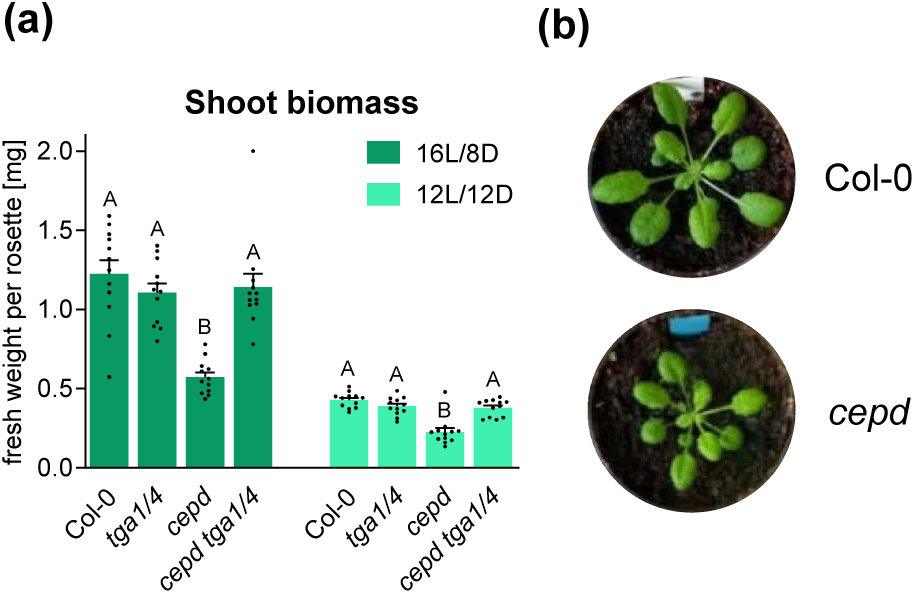
CEPDs counteract the repressive effect of TGA1/4 on rosette biomass formation. **(a)** Fresh weight of Col-0, *tga1 tga4*, *cepd* and *cepd tga1 tga4*. Plants were grown for 4 weeks under a 16 h light (L)/8 h dark (D) or a 12L/12D regime. Mean values of 12 biological replicates per genotype are shown. Error bars represent the standard error of the mean. Letters indicate statistically significant differences between the genotypes. Statistical analysis was performed by one-way ANOVA and Tukey’s post-test (*p* adj. < 0.05). **(b)** Picture of representative plants. Plants were grown for four weeks under a 12L/12D regime.

### CEPDs are required for the expression of genes related to transport of inorganic ions including nitrate

In order to identify genes that are under the control of CEPDs we first searched for conditions with strong differential expression levels of CEPDs. According to publicly available expression data (www.genevestigator.de; Supplemental Fig. 2a), *ROXY8* and *ROXY9* are higher expressed in seedlings cultivated for two days in liquid medium containing 0.15 mM NO_3_^-^ and 0.05 mM NH_4_^+^ (low nitrogen (LN); (Scheible et al., 2004)) as compared to controls grown in the presence of 3 mM NO_3_^-^, 1 mM NH_4_^+^ and 1 mM glutamine (full nutrition (FN) (Scheible et al., 2004)). In order to be able to investigate gene expression in shoots and roots separately, we grew seedlings on solid FN medium in square petri dishes. After seven days of growth under continuous light, seedlings were either transferred to LN or to FN media. After two further days, shoots and roots were harvested for RNA analysis.

As described for Noessen (Ohkubo et al., 2017; Ota et al., 2020), *CEPD* transcript levels were higher in shoots than in roots in Col-0 (Supplemental Fig. 2b). Upon transfer to medium with low N content, transcription of especially *ROXY8* and *ROXY9* increased in shoots, although to different degrees depending on the experiment (Supplemental Fig. 2b-d). *ROXY6* was induced only in one of the three experiments, while *ROXY7* expression was consistently independent of the N supply. TGA1/4 were required as transcriptional activators of *ROXY6*, *ROXY8* and *ROXY9* under FN, but not under LN conditions (Supplemental Fig. 2c). Expression of *ROXY6* was reduced in the *cepr1-3* mutant (Chapman et al., 2019) under FN and LN conditions while expression of *ROXY8* and *ROXY9* was elevated under both conditions (Supplemental Fig. 2d). This was in view of previously reported results on severely reduced expression of *ROXY6/CEPD1* and *ROXY9/CEPD2* in *cepr1-1* (Ohkubo et al., 2017) unexpected and might either be due to the different ecotypes used for the analyses or differences in the growth conditions.

*TGA1* transcript levels were higher in roots than in shoots while *TGA4* transcript levels were not different (Supplemental Fig. 3a). At the protein level, a band reacting specifically with the αTGA1 antibody was much more abundant in root-than in leaf extracts from soil-grown plants (Supplemental Fig. 3b). *TGA1* levels were only slightly increased upon growth on reduced N supply while *TGA4* transcript remained unaffected (Supplemental Fig. 3a).

RNAseq analysis was performed with RNA from roots of Col-0 and *cepd* subjected to LN conditions. Samples from four independent experiments with each sample containing roots from 50 plantlets grown on five separate plates were sent for sequencing. After Principal Component Analysis (PCA), we had to exclude one data set from Col-0 since the corresponding sample had been contaminated with leaf material. The remaining replicates revealed that 350 genes were less expressed and 212 genes were higher expressed in *cepd* compared to Col-0 (log_2_ FC >/< |1|, adjusted *p*-value (*p* adj) < 0.05, Supplemental Table 1).

Gene Ontology (GO) term analysis of the 350 genes that require CEPDs for expression unraveled an overrepresentation of genes involved in transport processes of inorganic ions including nitrate (Fig. 2a). This is consistent with the findings that CEPDs are responsible for nitrate acquisition in *Arabidopsis thaliana* (Ota et al., 2020; Ohkubo et al., 2021). Five of the ten most highly differentially expressed genes (log_2_ FC > 3.19) encode for transport proteins, including the two high-affinity nitrate transporters NRT2.2 and NRT2.4 (Zhuo *et al.,* 1999; Kiba *et al.,* 2012), the amino acid transporter UMAMIT35 (Zhao *et al.,* 2021), and the ammonium transporter AMT1-5 (Yuan *et al.,* 2007) (Fig. 2b). One gene encodes for NIN-LIKE PROTEIN 3 (NLP3), which is a nitrate-sensitive transcription factor (Liu *et al.,* 2022). The influence of CEPDs on expression of *NRT2.1*, which plays a major role in high-affinity NO_3_^−^ uptake in the root (Cerezo et al., 2001; Filleur et al., 2001), was less pronounced (log_2_ FC 2.04). The gene encoding phosphatase CEPH, which activates high-affinity nitrate uptake by dephosphorylating NRT2.1 (Ohkubo *et al.,* 2021), is among the top ten differentially regulated genes (log_2_ FC 3.22).

**Fig. 2.**
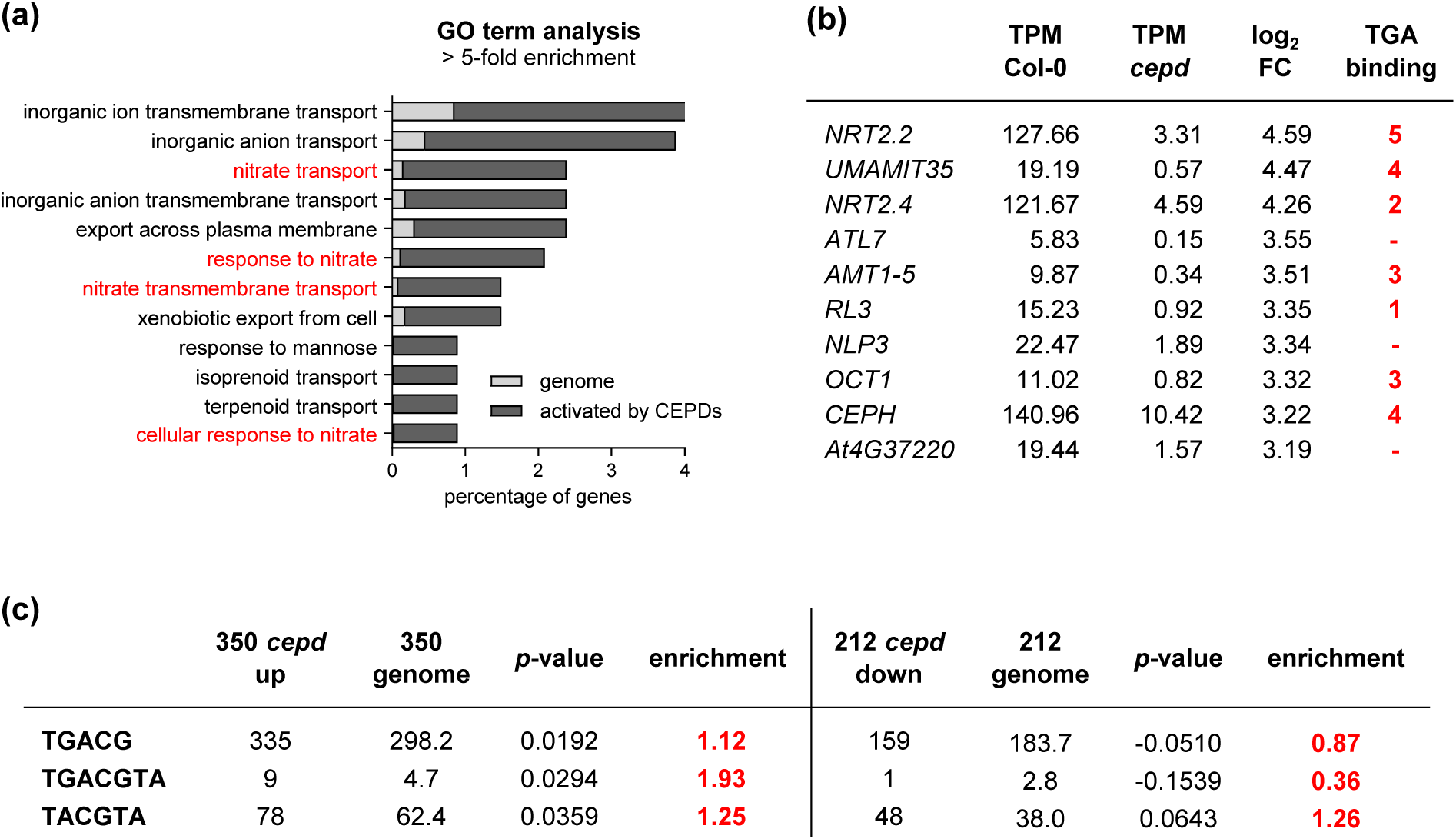
CEPDs are required for the expression of genes related to N acquisition. **(a)** Gene Ontology (GO) term analysis (biological processes) of 350 differentially expressed genes (log_2_ FC > 1, *p* adj. < 0.05) in roots of 9-day-old *cepd* seedlings compared to Col-0 after two days of cultivation on LN (low nitrogen) medium. GO terms related to nitrogen are displayed in red. Bars represent the percentage of genes found per GO term in the group of 350 genes (dark grey) and the percentage of genes representing the respective GO term found within the Arabidopsis genome (light grey). GO terms with > 5-fold enrichment against the genome are shown. Statistical analysis was performed using Fisher’s Exact test and False Discovery Rate (FDR) < 0.05. **(b)** List of the ten most highly differentially expressed genes. The number of TGA binding sites (TGACG and TACGTA) was counted in the region 2 kb upstream of the transcriptional start site. TPM: transcripts per million. **(c)** Motif Mapper *cis*-element analysis. Numbers of motifs in sequences 1 kb upstream of the transcriptional start site in 350 differentially expressed genes and in 350 genes randomly picked from the genome are indicated.

Finally, motif mapper analysis (Berendzen et al., 2012) documented enrichment of TGA factor binding motifs TGACG, TGACGTCA and TACGTA (Izawa et al., 1993; Wang et al., 2019) in the promoter regions of those genes that are positively regulated by CEPDs (Fig. 2c).

### CEPDs interfere with the repressive effect of TGA1 and TGA4 on expression of N starvation-induced genes

Next, we tested the expression pattern of the top three differentially regulated genes (Col-0 vs. *cepd*, log_2_ FC > 4.26) in roots of *tga1 tga4* and *cepd tga1 tga4* plants both under FN and LN conditions (Fig. 3). Upon transfer of plants to LN medium, *NRT2.2* was highly induced in Col-0, but not in *cepd*. Compromised expression in *cepd* was reverted to wild-type transcript levels in *cepd tga1 tga4*. Apparently, TGA1/4 strongly interfere with LN-induced *NRT2.2* expression and this repressive effect needs to be released by CEPDs while CEPD-TGA1/4-independent mechanisms confer strong up-regulation of the gene under N limiting conditions as compared to full N supply.

**Fig. 3.**
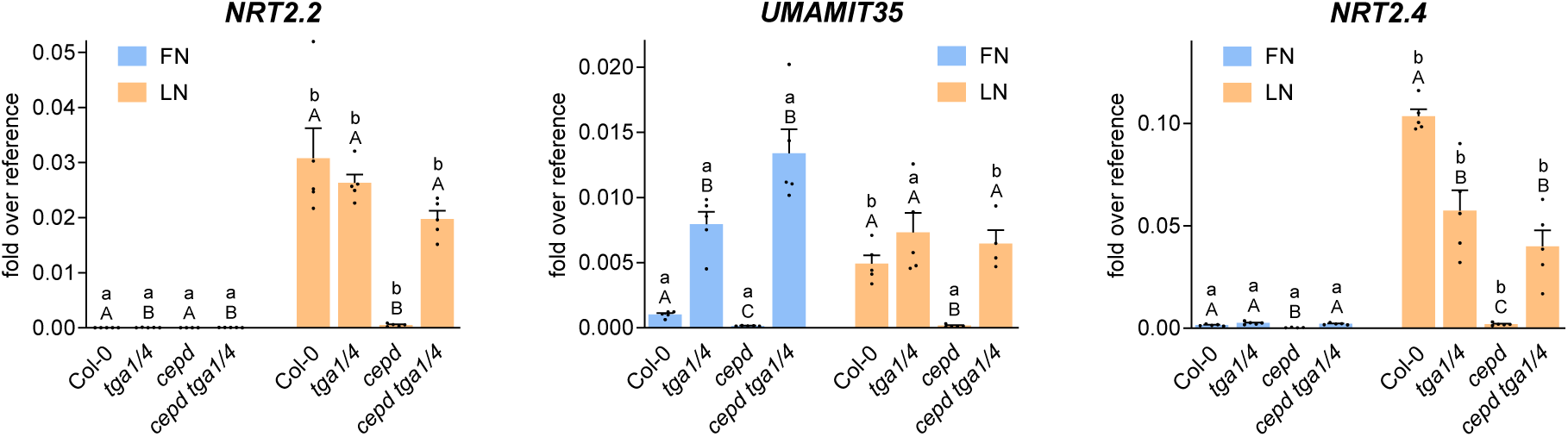
Impaired gene expression in *cepd* is due to the repressive effect of TGA1/4. 7-day old seedlings grown on full nitrogen (FN) medium under constant light (70 µmol photons s^-1^ m^-2^) were transferred to either FN or low nitrogen (LN) plates. Two days later, shoots and roots were collected for RNA isolation. Expression of the indicated genes was analyzed by qRT-PCR, *UBQ5* was used as a reference gene. Mean values of four to five biological replicates are shown, with one replicate originating from one plate with 10 plantlets. Error bars represent the standard error of the mean. Lowercase letters indicate statistically significant differences within the genotype between the treatments, uppercase letters indicate significant differences within treatment between the genotypes. Statistical analyses were performed with the logarithmic values by using two-way ANOVA and Bonferroni’s post-test (*p* adj. < 0.05).

In contrast to *NRT2.2*, *UMAMIT35* was higher expressed in *tga1 tga4* and *cepd tga1 tga4*, at least under FN conditions. Upon transfer of plants to LN, *UMAMIT35* expression was induced in Col-0 reaching similar levels as in *tga1 tga4*. This can be explained by LN-induced expression of *CEPDs* (Supplemental Fig. 2), leading to a maximal repressive effect on the negative activity of TGA1/4. In *cepd*, *UMAMIT35* expression was low due to the repressive activity of TGA1/4. Again, the *cepd tga1 tga4* hexuple mutant showed similar *UMAMIT35* expression levels as the *tga1 tga4* mutant. Unlike *NRT2.2*, *UMAMIT35* expression was not regulated by the N content in the medium in *cepd tga1 tga4*. Apparently, an N supply-independent activator leads to transcriptional activation. The regulation is installed by TGA1/4 and CEPDs, with TGA1/4 interfering with constitutive transcription and being de-activated under LN conditions by increased levels of CEPDs.

The regulation of *NRT2.4* was similar to *NRT2.2*. However, under LN conditions, when the repressive activity of TGA1/4 was released by CEPDs, TGA1/4 contributed to transcriptional activation.

All three genes were lower expressed in *cepr1-3* (Supplemental Fig. 4) although *ROXY8* and *ROXY9* transcript levels were elevated (Supplemental Fig. 2). It seems, that either ROXY6 or other gene products that are less abundant in *cepr1-3* are required for induction of *NRT2.2*, *UMAMIT35* and *NRT2.4*.

### CEPDs interfere with the expression of many genes related to photosynthesis

GO term analysis of the 212 genes that were more strongly expressed in *cepd* than in Col-0 unravelled an overrepresentation of genes related to photosynthesis (Supplemental Fig. 5a). The TGACG motif is depleted, but the TACGTA motif, which is bound *in vivo* by TGA1/4 (Wang et al., 2019), is slightly enriched (Fig. 2c). The ten most strongly differentially expressed genes contain TGA binding sites in their promoters (Supplemental Fig. 5b).

The most highly CEPD-repressed gene that we identified is *At4g39675* which encodes a 70 amino acid long hypothetical protein. The corresponding transcript levels were strongly down-regulated after transfer of plants from FN to LN media (Fig. 4). Under FN conditions, *At4g39675* was not affected by the CEPD-TGA1/4 regulatory module, suggesting that it was activated by a mechanism that was not repressed by CEPDs. However, under LN conditions, CEPDs repressed the activating function of TGA1/4.

**Fig. 4.**
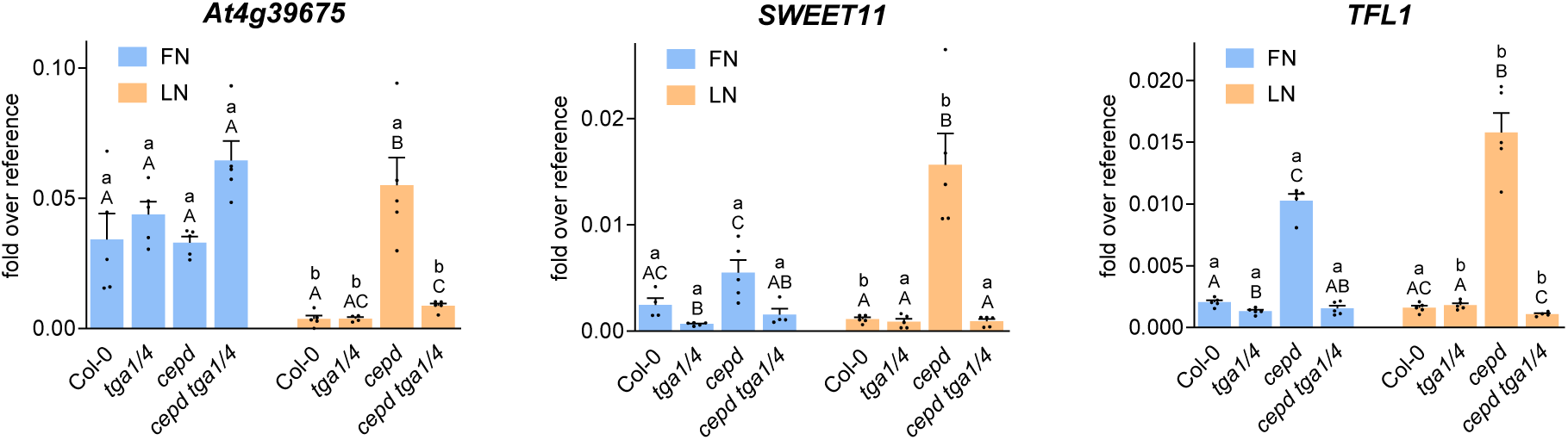
Enhanced expression in cepd is due to the activating effect of TGA1/4. 7-day old seedlings grown on full nitrogen (FN) medium were transferred to either FN or low nitrogen (LN) plates. Two days later, shoots and roots were collected for RNA isolation. Expression of the indicated genes was analyzed by qRT-PCR, *UBQ5* was used as a reference gene. Mean values of four to five biological replicates are shown, with one replicate originating from one plate with 10 plantlets. Error bars represent the standard error of the mean. Lowercase letters indicate statistically significant differences within the genotype between the treatments, uppercase letters indicate significant differences within treatment between the genotypes. Statistical analyses were performed by using two-way ANOVA and Bonferroni’s post-test (*p* adj. < 0.05).

Expression of a member of the SWEET sucrose efflux transporter family proteins (*SWEET11*) was also reduced upon transfer of plants from FN to LN, although not nearly as strongly as *At4g39675*. Here, the roles of TGA1/4 as CEPD-repressed activators can be observed under FN and LN conditions, especially in the *cepd* background. Down-regulation under LN conditions can be explained by an increased repressive effect of CEPDs on TGA1/4-mediated activation.

*TERMINAL FLOWER1* (*TFL1*) expression was strongly repressed by CEPDs both under FN and LN conditions. The activating effect of TGA1/4 can only be detected in the absence of CEPDs. Due to the strong repression of this gene by CEPDs already under FN conditions, LN conditions cannot further diminish gene expression.

All three genes were higher expressed in *cepr1-3* (Supplemental Fig. 6) although *ROXY8* and *ROXY9* transcript levels were elevated in this mutant (Supplemental Fig. 2). It seems, that either ROXY6 or other gene products that are less abundant in this mutant are required for interfering with transcriptional activation of *At4g39675, SWEET11* and *TFL1*.

### FN-induced putative repressive CC-type glutaredoxin-like proteins ROXY10-15 do not explain the negative activity of TGA1 and TGA4 on transcription

In view of published data that TGA1/4 can bind directly to TGACG motifs in the promoter regions of *NRT2.1* and *NRT2.2* (Alvarez et al., 2014) and considering the enrichment of TGA binding sites in promoters that require CEPDs for activation (Fig. 2c), we hypothesized that TGA1/4 mediate the repressive effect upon binding to these promoters. Since TGA1/4 have been described as direct transcriptional activators of defense, N dose- and nitrate-induced genes (Canales et al., 2017; Sun et al., 2018; Swift et al., 2020), we speculated that TGA1/4 might be associated with a co-repressor such as e.g. TOPLESS (Long et al., 2002) when regulating N starvation response genes. TOPLESS binds to an ALWL motif, which is present at the C terminus of all CC-type glutaredoxin-like proteins except ROXY20 and CEPDs (Uhrig et al., 2017). This ALWL motif is required for repression of PAN and TGA2 activity (Li et al., 2009; Zander et al., 2012). Since TGA1 and TGA4 can interact at least with ROXY18 and ROXY19 (Li et al., 2019), ROXYs encoding an ALWL motif might lead to the establishment of a repressive complex on TGA1/4 under FN conditions; this complex might be displaced by LN-induced CEPDs.

It has been described before that ALWL-containing CC-type glutaredoxin-like proteins *ROXY10,11,12,13,14,15* are more highly expressed in seedlings grown under FN than under LN conditions (www.genevestigator.de (Scheible et al., 2004), Supplemental Fig. 2a) and might therefore recruit TOPLESS to TGA1/4 especially under FN conditions. The tandem arrangement of *ROXYs 11-15* at one locus allowed deletion of all five genes by CRISPR/Cas9 genome editing (Supplemental Fig. 1). We also generated the *roxy10* mutant and crossed it with *roxy11-15*.

We observed a negative effect of ROXY10-15 on *UMAMIT35* expression, which was – however - not as pronounced as the negative effect of TGA1/4 (Supplemental Fig. 7). In addition, we tested two other LN-induced CEPD-dependent genes, namely *CLAVATA3-RELATED3* (*CLE3*) and *PEROXIDASE10* (*PER10*), which show pronounced repression by TGA1/4 under FN conditions (Fig. 5). However, ROXY10-15 did not influence expression of these genes. We therefore reject our hypothesis that ROXY10-15 are responsible for the strong repressive effect of TGA1/4 under FN conditions. Conspicuously, transcription of *CLE3* and *PER10* was reduced in *tga1 tga4* upon transfer from FN to LN, unravelling that TGA1/4 mask the effect of other regulatory elements influencing expression in an N supply-dependent manner.

**Fig. 5.**
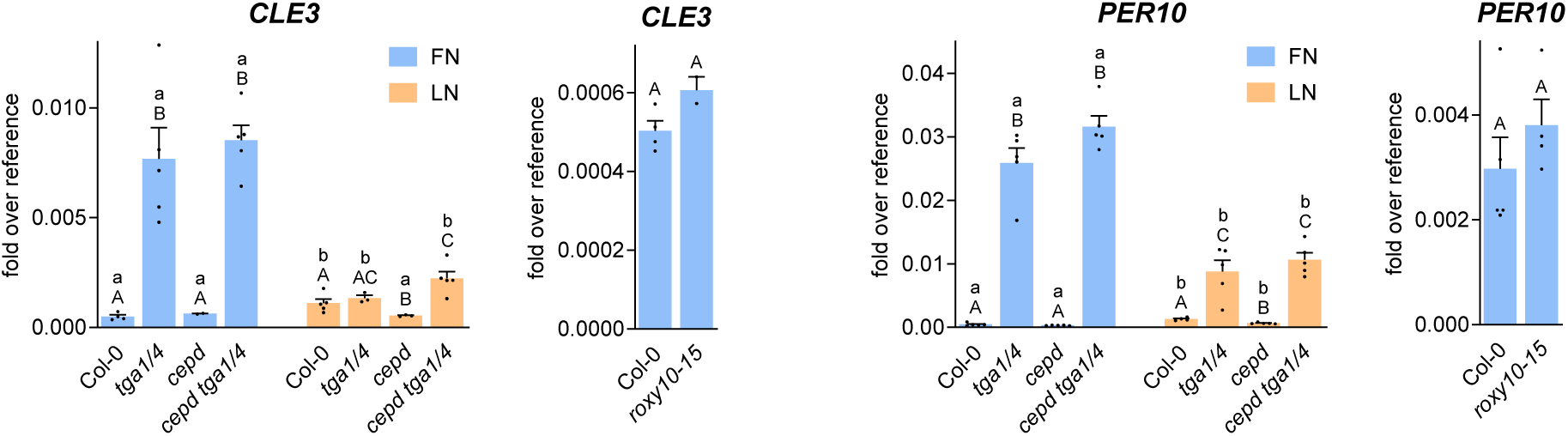
ALWL motif-containing ROXYs 10-15 are not responsible for repression of CEPD-activated genes. 7-day-old seedlings grown on full nitrogen (FN) medium were transferred to either FN or low nitrogen (LN) plates. Two days later, roots were collected for RNA isolation. Expression of the indicated genes was analyzed by qRT-PCR, *UBQ5* was used as a reference gene. Mean values of four to five biological replicates are shown with one replicate originating from one plate with 10 plantlets. Error bars represent the standard error of the mean. Lowercase letters indicate statistically significant differences within the genotype between the treatments, uppercase letters indicate significant differences within treatment between the genotypes. Statistical analyses were performed with the logarithmic values by using two-way ANOVA and Bonferroni’s post-test (*p* adj. < 0.05) or unpaired t-test (*p* < 0.05) for Col-0/*roxy10-15* comparison.

### CEPDs interfere with the repressive function of TGA1 and TGA4 on transport of nitrate to the shoot

As shown in Fig.1, loss of CEPDs leads to reduced rosette fresh weight in plants grown on fertilized soil re-enforcing the notion that CEPDs are not only important under N starvation but also under N replete conditions (Ota et al., 2020). Consistently, *cepd* seedlings accumulated less shoot biomass, irrespective of whether they were grown on 10 or 1 mM NO_3_^-^ (Fig. 6a). Shoot biomass did not vary significantly between Col-0 and *tga1 tga4* indicating that the repressive effect of TGA1/4 on shoot growth is completely counter-acted by CEPDs although their expression levels – shown exemplarily for *ROXY9* - are lower at 10 mM NO_3_^-^ than at 1 mM NO_3_^-^ (Supplemental Fig. 8a). Fresh weight of *tga1 tga4* roots was reduced at 10 mM NO_3_^-^ (Fig. 6a), indicating that TGA1/4 have root growth-activating functions that are not or not fully repressed by CEPDs. The *cepd* mutant also accumulated less root fresh weight. No additive effect was observed in the *cepd tga1 tga4* mutant. Differences in root fresh weight between the genotypes were not significant at 1 mM NO_3_^-^.

**Fig. 6.**
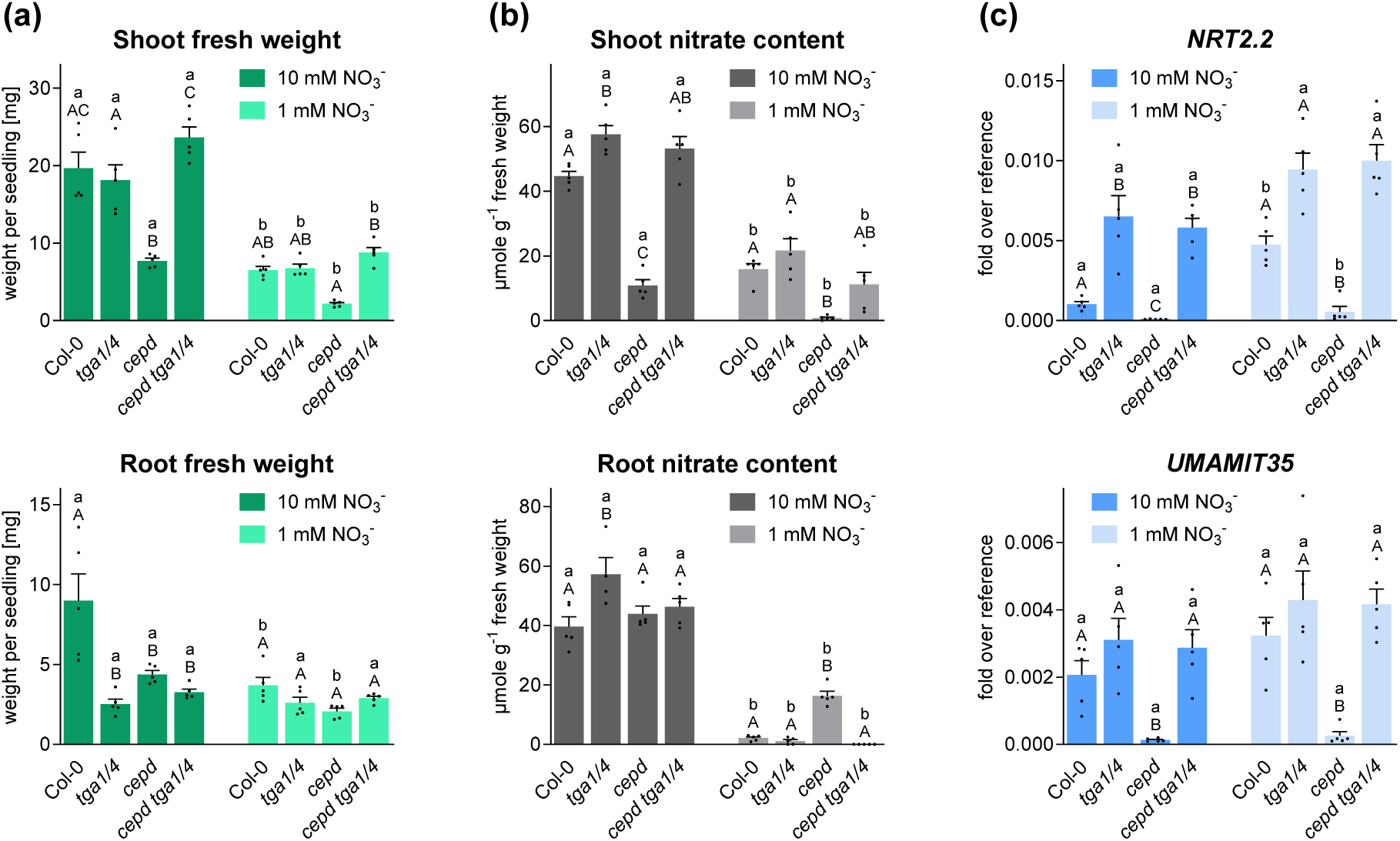
Abnormal nitrate allocation and shoot biomass in *cepd* is reverted in *cepd tga1 tga4*. Wild-type (Col-0), *tga1 tga4, cepd* and *cepd tga1 tga4* plants were grown on medium containing 1 or 10 mM NO_3_^-^ under constant light (70 µmol photons s^-1^ m^-2^). After 21 days, shoots and roots were collected separately for determination of biomass **(a)**, nitrate content **(b)** and transcript levels **(c)**. Mean values of five biological replicates are shown, with one replicate originating from one plate with 10 plantlets. Error bars represent the standard error of the mean. Lowercase letters indicate statistically significant differences within the genotype between the treatments, uppercase letters indicate significant differences within treatment between the genotypes. Statistical analyses were performed with linear (a, b) or logarithmic values (c) by using two-way ANOVA and Bonferroni’s post-test (*p* adj. < 0.05).

Next, we measured the nitrate content in shoots and roots of Col-0, *tga1 tga4*, *cepd* and *cepd tga1 tga4* (Fig. 6b). Col-0 wild-type plants accumulated 2.8-fold more nitrate in the shoot when grown on 10 mM NO_3_^-^ as compared to 1 mM NO_3_^-^, while the difference in root nitrate content was 19-fold, indicating preferential nitrate transport to the shoot. When compared to Col-0, the shoot nitrate content of *cepd* was reduced by a factor of four at 10 mM NO_3_^-^ and 21-fold at 1 mM NO_3_-. Conspicuously, the root nitrate content was not altered in *cepd* at 10 mM NO_3_^-^ and was even 8-fold higher at 1 mM NO_3_^-^ . The nitrate content of total seedlings grown on 1 mM NO_3_^-^ was 10.8 µmol/g FW in Col-0 and 8.1 µmol/g FW in the *cepd* mutant. This underlines the notion that especially root to shoot allocation of nitrate is impaired in *cepd*, while uptake is less severely reduced. Elevated root nitrate content at the expense of shoot nitrate content in *cepd* was also observed when plants were grown on 3 mM NO_3_^-^ (Supplemental Fig. 9). The *cepd* phenotype with respect to nitrate content in shoots and roots was completely suppressed by *tga1 tga4* alleles.

At the level of gene expression, we again observed that expression of genes related to nitrate uptake under N limiting conditions like *NRT2.1*, *NRT2.2*, *NRT2.4* and *CEPH* was reduced in *cepd* (Fig. 6c, Supplemental Fig. 8b). Dual affinity transporters like *NRT1.1*, which were not repressed by TGA1/4 in the absence of CEPDs (Supplemental Fig. 8b), may account for the uptake of NO_3_^-^ that we observed in *cepd* even at 1 mM NO_3_^-^ (Fig. 6b) (Liu and Tsay, 2003). Since nitrate root to shoot allocation was more affected than nitrate uptake, we tested expression of *NRT1.5*, which encodes a low-affinity nitrate transporter that participates in nitrate loading of the root xylem (Lin et al., 2008). However, *NRT1.5* expression was not as stringently affected in *cepd* as expression of *NRT2.2* and *NRT2.4* (Supplemental Fig. 8b). Reduced expression of all analyzed genes in *cepd* was completely suppressed by *tga1 tga4* alleles (Fig. 6c, Supplemental Fig. 8b).

In contrast to what we had observed in plants cultivated on FN medium (Fig. 3), *NRT2.2* expression was higher in *tga1 tga4* than in Col-0 during growth on 10 mM NO_3_^-^ (Fig. 6c). Apparently, ammonium and glutamine in the FN medium repress *NRT2.2* expression in Col-0 and *tga1 tga4* (Supplemental Fig. 10a). *NRT2.2* expression was induced when reducing the nitrate supply from 10 to 1 mM presumably because of the almost complete release of the repressive TGA1/4 function by increased CEPD levels (Fig. 6c, Supplemental Fig. 8a). Other CEPD-regulated genes like *NRT2.1*, *NRT2.4* and *NRT1.5* showed similar expression levels in Col-0 and *tga1 tga4* on 1 mM and 10 mM NO_3_^-^ (Supplemental Fig. 8b) suggesting that the negative effect of TGA1/4 was fully lifted by CEPDs independent of the amount of nitrate. Expression levels of *NRT2.1*, *NRT2.4* and *NRT1.5* in Col-0 were similar under sufficient (10 mM N) and limiting (1 mM N) N supply, which is – at least in the case of *NRT2.4* - different from the situation on FN (6 mM N) and LN (0.2 mM N) medium (Fig. 3). Repression of *NRT2.4* by ammonium and glutamine accounts for very low basal transcript levels in plants grown on FN.

Induction of *UMAMIT35* expression upon transfer of plants from FN to LN had been due to increased *CEPD* expression and thus complete release of TGA1/4-mediated repression (Fig. 3). When plants were grown on 10 mM NO_3_^-^ as compared to 1 mM NO_3_^-^ (Fig. 6c), gene expression was not affected by the N supply. This correlates with the finding that *UMAMIT35* expression was not elevated in *tga1 tga4* indicating that CEPDs completely interfere with TGA1/4 activity when plants face 1 or 10 mM NO_3_^-^. Whether compromised CEPD activity in plants cultivated on FN is due to lower expression levels of CEPDs or other mechanisms, has remained an open question (Supplemental Fig. 10b).

It is concluded that the efficiency of the repressive effect of CEPDs on TGA1/4-mediated repression depends on the promoter context and that glutamine/ammonium in the medium control promoters through CEPD-dependent (*UMAMIT35*) and CEPD-independent (*NRT2.2, NRT2.4*) mechanisms.

### Reactive cysteines in TGA1 are not involved in the regulation of CEPD-dependent genes

We have shown previously that TGA1/4 and ROXY9 interact in yeast and in transiently transformed protoplasts (Li et al., 2019), implicating that TGA1/4 activity in roots is regulated by CEPDs through protein-protein interactions. Since TGA1/4 contain redox active cysteines (Despres et al., 2003; Lindermayr et al., 2010) and considering the high sequence similarity of CEPDs to glutaredoxins, it seemed plausible that CEPDs affect TGA1/4 activity by regulating their redox state. Therefore, we analyzed the induction of N starvation response genes in previously described *tga1 tga4* complementation lines which express equivalent amounts of either TGA1 or a TGA1 variant in which the four cysteines of the protein were mutated (Budimir et al., 2021) (Supplemental Fig. 3b). Since the latter construct mimics the reduced state, we had called it TGA1_red_. The *TGA1* constructs contain the endogenous promoter, 5’ and 3’ UTRs, all introns and an HA tag at the N terminus.

In shoots of soil-grown plants, *ROXY9* expression was lower in *tga1 tga4* than in Col-0, while *ROXY13* expression was higher. HA-TGA1 and HA-TGA1_red_ complemented expression of these genes to a similar extent (Fig. 7a). Likewise, HA-TGA1 and HA-TGA1_red_ were almost equally efficient in repressing *UMAMIT35* expression under FN conditions (Fig. 7b), suggesting that TGA1 can function as a repressor in its pseudo-reduced form. If inactivation of TGA1/4 by LN-induced CEPDs would require oxidation of cysteine residues, *UMAMIT35* induction would be impaired in the transgenic line expressing the mutated protein. However, induction was possible, indicating that the repressive activity of TGA1 is not lifted by changes in its redox state. *PER71* belongs to the group of genes that is activated by TGA1/4 and repressed by CEPDs, with increased CEPD activity under LN conditions leading to weaker expression levels in plants grown on LN (Supplemental Fig. 11). HA-TGA1 and HA-TGA1_red_ activated *PER71* to similar levels, and expression was reduced in a similar manner in the two complementation lines under LN conditions. Thus, CEPD-mediated inactivation of TGA1 as a transcriptional activator is not due to redox modulation of cysteines in TGA1.

**Fig. 7.**
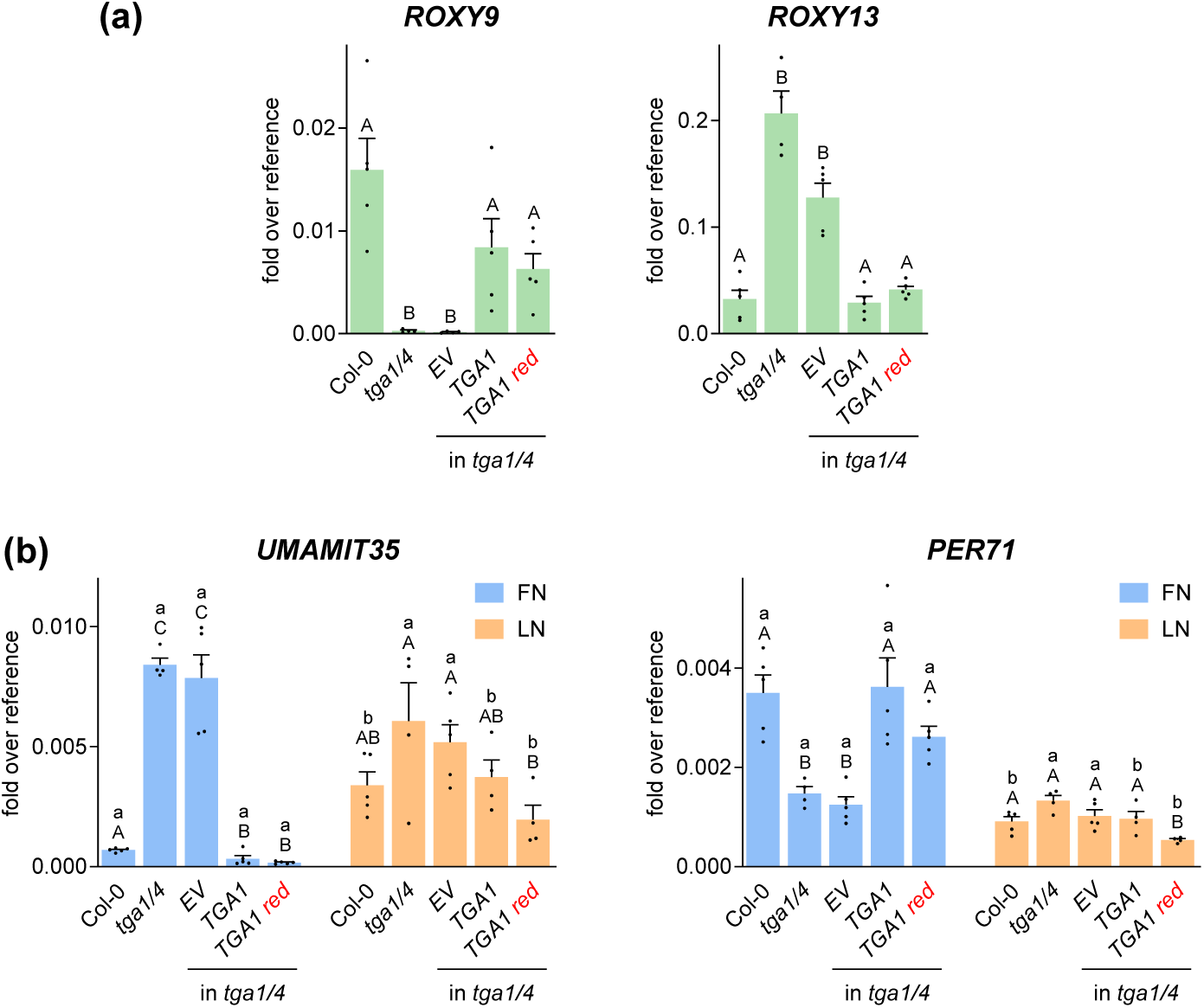
Regulation of TGA1 activity is not due to modification of its redox state. **(a)** qRT-PCR analysis of *ROXY9* and *ROXY13* transcript levels in Col-0, *tga1 tga4* and respective complementation lines. RNA is from leaves of 4-week-old soil-grown *A. thaliana* plants. *UBQ5* was used as a reference gene. Mean values of samples from three to six plants are shown. Error bars represent the standard error of the mean. Statistical analyses were performed with the logarithmic values using one-way ANOVA and Tukey’s post-test (*p* adj. < 0.05). **(b)** qRT-PCR analysis of *UMAMIT35 and PER71* in 9-day-old *A. thaliana* seedlings. Plants grown on full nitrogen (FN) plates were transferred to either FN or low nitrogen (LN) plates. Two days later, RNA was isolated from roots. *UBQ5* was used as a reference gene. Mean values of four to five biological replicates are shown with one replicate originating from one plate with 10 plantlets. Error bars represent the standard error of the mean. Lowercase letters indicate statistically significant differences within the genotype between the treatments, uppercase letters indicate significant differences within treatment between the genotypes. Statistical analyses were performed with the logarithmic values by using two-way ANOVA and Bonferroni’s post test (*p* adj. < 0.05). EV: empty vector control

### CEPDs modulate the activating function of TGA1 and TGA4 on hyponastic growth

Next, we aimed to explore whether CEPDs are involved in TGA1/4-regulated processes in the shoot. Under unfavourable conditions such as submergence, high temperatures, darkness (D) or low light (LL) (Polko et al., 2011), plants bend their leaves upwards (hyponasty) by elongation of the abaxial epidermal cells of the petioles (Kupers et al., 2023). To reverse this process (epinasty), the adaxial cells of the petioles elongate (Polko et al., 2012). *tga1 tga4* and plants ectopically expressing HA-ROXY9 are impaired in hyponastic growth underpinning the concept that ROXY9 is a repressor of TGA1/4 function (Li et al., 2019). Since *ROXY8* and *ROXY9* transcript levels are up-regulated upon transfer of plants from LL to normal light (NL), we had suggested that ROXY8 and ROXY9 counteract the hyponasty-activating function of TGA1/4 to allow downward movement of leaves (Li et al., 2019).

In order to provide loss of function evidence for a potentially regulatory role of CEPDs on hyponastic growth, Col-0, *tga1 tga4*, *cepd* and *cepd tga1 tga4* plants, grown for four weeks under a 12h NL (100-120 µmol photons m^2^ s^-1^)/12h D regime, were transferred to LL (15-20 µmol photons m^2^ s^-1^) for 10.5 hours (Fig. 8a). After the following dark period, plants were kept for another four hours under LL and petiole angles were measured. As described before, the LL-induced hyponasty was reduced in *tga1 tga4* (Li et al., 2019) (Fig. 8b). Petioles of *cepd* plants that had been kept in parallel under the NL/D regime were in a slightly more upward position compared to Col-0. Though starting already at a higher petiole angle, *cepd* plants responded to the LL stimulus, indicating that activation of hyponasty is not (only) due to the release of the repressive activity of CEPDs on the hyponasty-promoting function of TGA1/4. Unexpectedly, petiole angles of *cepd tga1 tga4* responded to LL suggesting the existence of TGA1/4-independent hyponasty-promoting factors that are repressed by CEPDs. Taken together, these results demonstrate that CEPDs dampen TGA1/4-dependent and TGA1/4-independent hyponasty.

**Fig 8.**
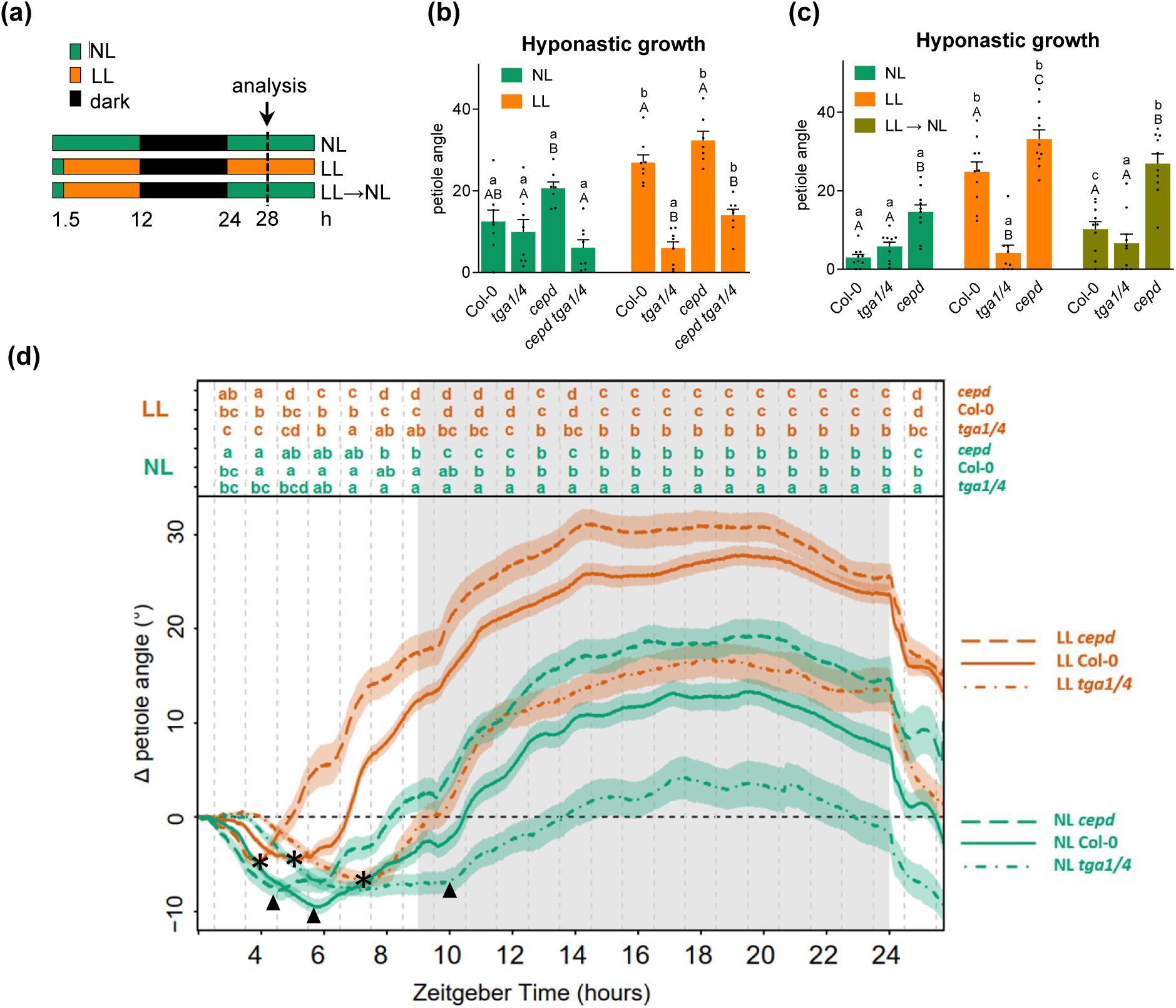
CEPDs dampen hyponastic growth-promoting functions. **(a)** Scheme of the light conditions. Wild-type Col-0, *tga1 tga4*, *cepd* and *cepd tga1 tga4* plants were grown under 12 h normal light (NL; 100-120 µmol photons m^−2^ s^−1^)/12 h D regime. Plants were transferred to low light (LL; 15-20 µmol photons m^−2^ s^−1^) for 10.5 h, followed by a 12 h dark period. On the next day, plants were cultivated for 4 further hours in LL (b) or NL (c). The leaf angle of the 8^th^ leaf was measured with the ImageJ software. Bars represent the average ± SEM of eight plants per genotype. Error bars represent the standard error of the mean. Lowercase letters indicate statistically significant differences within a genotype between different light conditions, uppercase letters indicate significant differences between genotypes subjected to the same light conditions. Statistical analysis was performed by using two-way ANOVA and Bonferroni’s post-test (*p* adj. < 0.05). (d) Time-resolved tracking of hyponastic growth. Eight plants per genotype were grown under an NL/D cycle. Petiole angles of the different genotypes were set to 0 at ZT=2 and the relative angle change was measured every minute for 24 h. At ZT=2, plants were transferred to LL (approximately 25 µmol photons m^−2^ s^−1^). The grey area indicates the dark period. Letters indicate *p* adj. < 0.05, calculated per every hour using one-way ANOVA and Tukey post-test. Stars and triangles indicate the start of hyponastic growth under NL and LL, respectively.

When plants were kept for 10.5 hours under LL before the 12 hour dark period and cultivated subsequently under NL for four further hours (Fig. 8c), the LL-induced increase in petiole angle was reverted in Col-0. This re-orientation was less pronounced in *cepd*, which is consistent with our hypothesis that CEPDs might be required to dampen the hyponasty-promoting activity of TGA1/4 when plants are transferred from LL to NL.

Time-resolved tracking of petiole angles of plants cultivated in a 9h NL/15h D or 9h LL/15h D regime extended the results of the end-point measurements. Under the normal NL/D rhythm, Col-0 plants move their petioles in a diurnal pattern (Oskam et al., 2023) with petioles moving down during the first six hours of the light period after which they move up again three hours before the expected night period (Fig. 8d; Supplemental Fig. 12). Hyponastic growth continues during the first five hours (until Z=14) of the night, after which it levels off and epinastic growth starts at around ZT=19. Downward movement strongly increases upon the onset of the light period. Since the amplitude of the diurnal hyponastic response is affected in *tga1 tga4* and *cepd*, petiole angles were already different at the start of the tracking period. Therefore, the petiole angles of the three genotypes were all set to 0 at 2 hours (ZT=2) after the end of the dark period and the relative change in the following 24 hours was monitored.

During the NL/D regime, petiole angles of *tga1 tga4* were smaller than those of Col-0 especially during the dark period with this difference starting to become significant roughly one hour after the onset of the dark period (ZT=10 to ZT=24) (Fig. 8d). In *cepd*, petiole angles were significantly larger during the period of hyponastic growth in the first half of the dark period between ZT=9 and ZT=14. Differences were partially due to the pre-ponement of the time point when growth switched from epinasty to hyponasty in *cepd* and its post-ponement in *tga1 tga4.* This observation is consistent with the notion that CEPDs interfere with TGA1/4 activity to allow complete downward movement after a period of upward movement (Fig. 8b). CEPDs did not influence the response in *tga1 tga4* (Supplemental Fig. 13), supporting the notion that CEPDs act by dampening TGA1/4 activity.

Under the LL/D regime, differences between Col-0, *tga1 tga4* and *cepd* became already visible during the five ours preceding the dark period. Again, the time point where growth switched from epinasty to hyponasty was pre-poned in *cepd* and postponed in *tga1 tga4* (Fig. 8d). When we compared *tga1 tga4* with *cepd tga1 tga4* (Supplemental Fig. 11), no significant change was observed between *cepd* and *cepd tga1 tga4* during the LL period confirming that CEPDs influence this phenotype only in the presence of TGA1/4. However, hyponastic growth during the dark after the LL period was partially restored in *cepd tga1 tga4* (Supplemental Fig. 13). This result is consistent with the findings obtained from the end-point measurements (Fig. 8b) and indicates that CEPDs repress - in the absence of TGA1/4 - other hyponastic growth-promoting factors that show enhanced activity in the dark after plants have experienced a LL light period.

### CEPDs only marginally interfere with the activating function of TGA1 and TGA4 on defense gene expression

TGA1/4 have been shown to be important for the expression of genes that are required for the establishment of the immune program systemic acquired resistance (SAR) (Sun et al., 2018; Nair et al., 2021). This program induces a state of alertness in systemic leaves after infection of local leaves with (hemi-)biotrophic pathogens (Durrant and Dong, 2004; Kachroo and Kachroo, 2020). Upon infection of SAR leaves, the elicited defense reaction is faster and stronger than upon infection of naive leaves. An important marker gene for this defense response is *FLAVIN-DEPENDENT-MONOOXYGENASE 1* (*FMO1, AT1G19250*). FMO1 catalyzes the hydroxylation of pipecolic acid (Pip) to yield the active signaling molecule N-hydroxy pipecolic acid (NHP), which is crucial for SAR establishment (Chen et al., 2018; Hartmann et al., 2018). *FMO1* expression is strongly impaired in the *tga1 tga4* mutant (Yildiz et al., 2023). To test, whether CEPDs dampen TGA1/4 activity during defense responses, expression of *FMO1* was analyzed in *cepd* mutants under SAR conditions (Fig. 9). Plants were either infiltrated with *Pseudomonas syringae* pv. *maculicola* (*Psm*) ES4326 or with a mock solution. After 48 hours, systemic leaves were infected with *Psm*. While *FMO1* expression was severely compromised in the *tga1 tga4* mutant, it was slightly but significantly enhanced in the *cepd* mutant, suggesting that CEPDs dampen TGA1/4 activity in the context of defense responses against *Psm* ES4326.

**Fig. 9.**
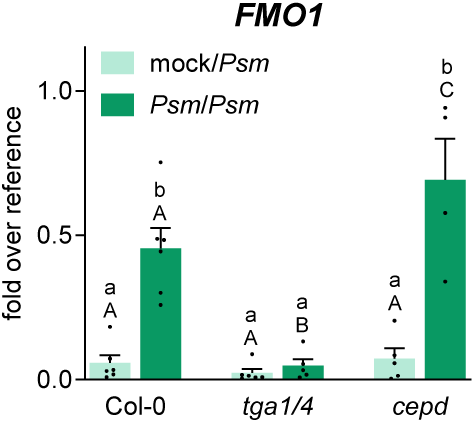
Induction of *FMO1* during SAR requires TGA1/4, but is only slightly repressed by CEPDs. RT-qPCR analysis of *FMO1* transcript levels in wild-type (Col-0), *tga1 tga4 and cepd* plants. Three leaves of 4.5-week-old soil-grown plants were infiltrated with either 10 mM MgCl_2_ (mock) or *Pseudomonas syringae* pv*. maculicola* (*Psm*) ES4326. Two days later, three systemic leaves were infiltrated with *Psm* and this tissue was harvested for RNA exctractiion at eight hours after infection. Expression of *FMO1* was analyzed by qRT-PCR, *UBQ5* was used as a reference gene. Mean values of four to six biological replicates are shown. Error bars represent the standard error of the mean. Lowercase letters indicate statistically significant differences within a genotype between different treatments, uppercase letters indicate significant differences between genotypes subjected to the same treatments. Statistical analysis was performed by using two-way ANOVA and Bonferroni’s post-test (*p* adj. < 0.05). Two independent experiments were performed with similar results.

## Discussion

CEPDs have been identified as N starvation-induced mobile shoot to root signals that are required for the transcriptional activation of genes related to N uptake and transport (Ohkubo et al., 2017; Ota et al., 2020). Here we show that CEPDs act by efficiently interfering with the strong repressive effect of transcription factors TGA1/4 (Fig. 10). CEPDs also repress TGA1/4-mediated transcriptional activation. The CEPD-TGA1/4 antagonism is less pronounced on shoot-borne TGA1/4 functions like activation of hyponasty or defense responses. CEPD-mediated regulation of TGA1/4 activity does not involve redox modification of critical cysteines in TGA1.

**Fig. 10.**
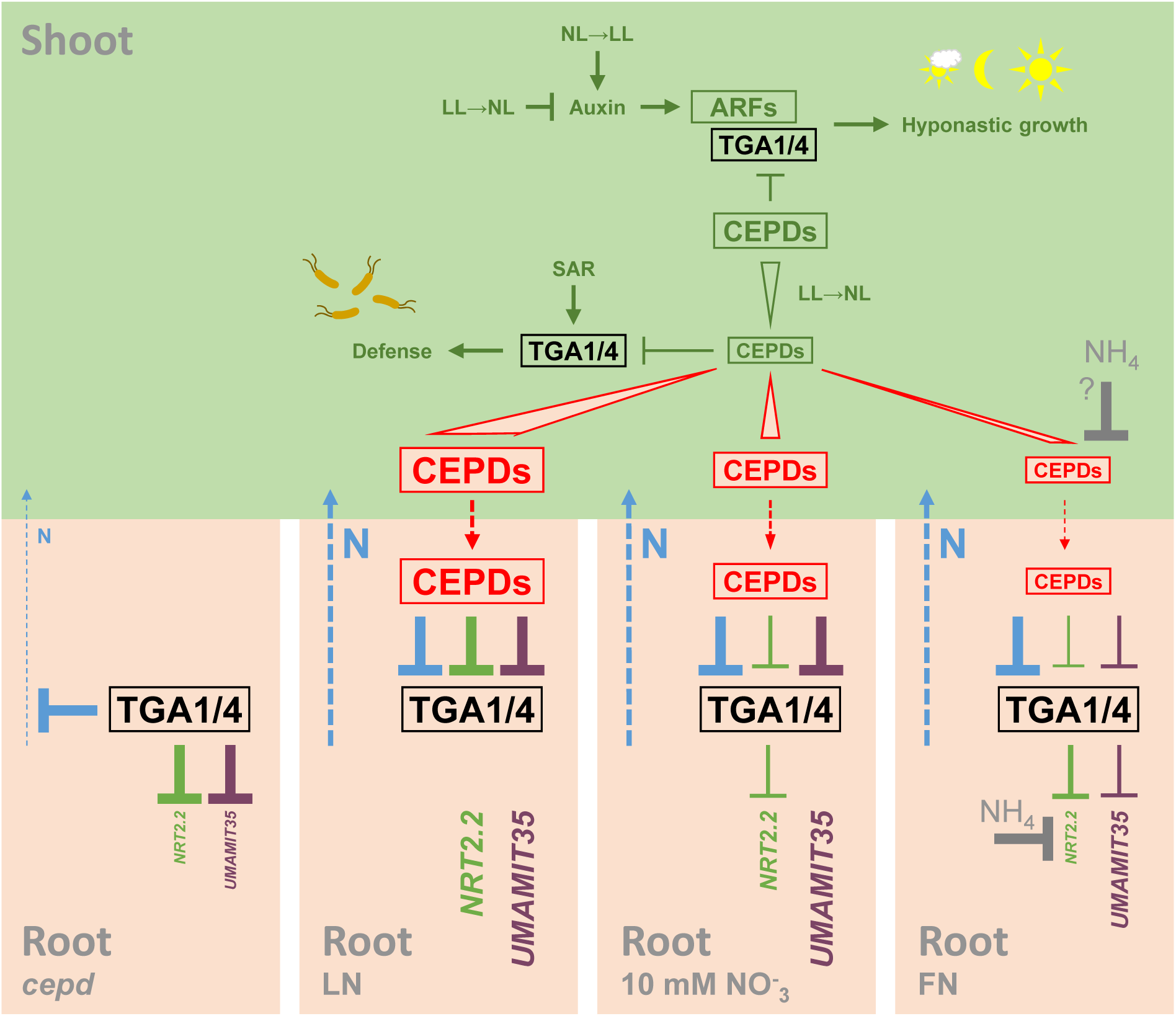
Regulation of TGA1/4 activities by CEPDs. In the shoot, TGA1/4 are important for defense responses in the context of systemic acquired resistance (SAR) and auxin-dependent hyponastic growth. These functions are dampened by CEPDs, which increase in the shoot upon transfer of low light (LL)-treated plants to normal light (NL) or under N starvation conditions. In *cepd* roots, genes involved in N translocation to the shoot and other genes like *NRT2.2* and *UMAMIT35* are strongly repressed by TGA1/4 irrespective of the N supply. Under low N supply (LN), CEPDs fully interfere with TGA1/4 activity allowing expression of the indicated marker genes *NRT2.2* and *UMAMIT35* and a yet unknown gene required for translocation of N to the shoot (blue). Under sufficient N supply being provided by 10 mM nitrate, *CEPD* expression in the shoot is lowered. Still, these levels are sufficient to fully interfere with the repressive activity of TGA1/4 activity on N translocation and *UMAMIT35* expression. Repression of *NRT2.2* is not fully lifted. Under full nutrition (FN), ammonium/glutamine repress *NRT2.2* independently of CEPDs and interfere with CEPD activity on the repressive action of TGA1/4 on *UMAMIT35*. Broken lines indicate translocation from shoot to root or vice versa, font sizes of *NRT2.2* and *UMAMIT35* represent relative expression levels, ARFs are auxin response factors mediating hyponastic growth.

### CEPDs act upstream of TGA1 and TGA4

Epistasis analysis is a genetic tool that unravels whether proteins act in one pathway and defines their hierarchy. Here, we observed that *cepd* phenotypes like reduced shoot weight (Figs. 1 and 6), altered nitrate content (Fig. 6) and impaired N starvation-induced gene expression (Figs. 3 and 6) depend on the presence of transcription factors TGA1/4 documenting that *tga1 tga4* alleles are epistatic to *cepd*. Similar studies have been performed for ROXY1 and TGA transcription factor PERIANTHIA (PAN) (Li et al., 2009). The *roxy1* mutant is characterized by in average only 2.5 instead of 4.0 petals characteristic for Arabidopsis wild-type flowers, while *pan* flowers have five petals (Chuang et al., 1999). Importantly, the *roxy1 pan* mutant has a *pan*-like phenotype with pentameric whirls. This indicates that the *roxy1* phenotype is due to a repressive function of PAN on the initiation of petal primordia, which is counteracted by appropriate amounts of ROXY1. Likewise, antagonistic roles of the three redundant CC-type glutaredoxin-like proteins MSCA1/ZmGRX2/ZmGRX5 and TGA factor FASCIATED EAR (FEA) 4 have been observed in the shoot apical meristem of maize: while the shoot apical meristems of *msca1 grx2 grx5* are smaller than wild-type meristems, those of *fea4* and *msca1 grx2 grx5 fea4* are larger. In contrast to *pan* (Chuang et al., 1999) and *fea4* (Pautler et al., 2015), which both have aberrant meristem phenotypes, the *tga1 tga4* mutant is similar to the wild-type when it comes to N acquisition and biomass production. This indicates that CEPDs completely interfere with the negative effect of TGA1/4 on processes regulating these phenotypes. If ROXY1 would completely suppress PAN activity, Arabidopsis plants would have pentameric flowers and if MSCA1/ZmGRX2/ZmGRX5 would completely suppress FEA4, maize meristems would be larger.

### TGA1 and TGA4 are CEPD-inactivated repressors of N uptake and transport genes

The strong negative effect of TGA1/TGA4 on growth and N acquisition processes has so far been overseen because it becomes evident only in the absence of CEPDs (Figs. 1 and 6). The *cepd* phenotype relates with altered gene expression in the root, where TGA1/4 function as repressors of N starvation-induced genes encoding high affinity nitrate transporters (NRT2.1, NRT2.2, NRT2.4) and the NRT2.1-regulating phosphatase CEPH (Fig. 3 and Supplemental Fig. 8). Over-representation of TGA binding sites in CEPD-activated promoters (Fig. 2b, c) and previous chromatin immunoprecipitation (ChIP) experiments documenting *in vivo* TGA1/4 binding to the promoter regions of *NRT2.1* and *NRT2.2* (Alvarez et al., 2014) suggest that TGA1/4 repress these genes upon binding to their target sites in these promoters. Since TGA1/4 have been described as transcriptional activators of defense, N dose- and nitrate responsive genes (Canales et al., 2017; Sun et al., 2018; Swift et al., 2020), we postulate that TGA1/4 recruit transcriptional co-repressors. Inactivation of this repressing complex by CEPDs allows expression of target promoters. Depending on the environmental conditions, TGA1/4 can then positively contribute to expression of *NRT2.1* and *NRT2.2*, as observed upon addition of nitrate to N starved roots (Alvarez et al., 2014). This notion is supported by our observation of a twofold activating effect of TGA1/4 on *NRT2.4* expression when plants were grown on LN medium (Fig. 3).

### Nitrate allocation to the shoot rather than nitrate uptake is strongly repressed by TGA1 and TGA4 in *cepd*

Under N limiting conditions, i.e. upon growth on 1 mM NO_3_^-^, *cepd* roots accumulated 8-fold higher nitrate levels than Col-0 roots, although expression of the major high affinity nitrate transporters *NRT2.1* (Cerezo et al., 2001; Filleur et al., 2001) and the corresponding activating phosphatase *CEPH* (Ohkubo *et al.,* 2021) was reduced by a factor of 2.7 and 3.3, respectively (Fig. 6, Supplemental Fig. 8). Expression of *NRT2.2* and *NRT2.4* was lowered by a factor of 9 and 29, respectively. However, the nitrate content of the shoot was reduced over 40-fold. Apparently, allocation of nitrate to the shoot rather than nitrate uptake is severely impaired in *cepd*. Consistently, a slightly increased root nitrate content under conditions where the shoot nitrate content was already reduced has been observed before in *cepdl2* (Noessen; corresponding to *roxy8* in Col-0) (Ota et al., 2020). With respect to the root to shoot allocation mechanism, only nitrate/potassium transporter NRT1.5 has been identified to facilitate excretion of nitrate from pericycle cells to the xylem but analysis of knock-out alleles demonstrated that other mechanisms have to be postulated as well (Lin et al., 2008). It remains to be explored whether the 2.2-fold reduction of *NRT1.5* mRNA levels in *cepd* (Supplemental Fig. 8) or low expression of other TGA1/4-repressed genes cause the strong allocation deficiency in *cepd*.

The deficiency in nitrate allocation to the shoot in *cepd* was only partially overcome upon growth on 10 mM NO_3_^-^ with shoot nitrate levels still being reduced by a factor of four. Ota *et al*. have reported a >50-fold reduction of shoot nitrate content when growing the *cepd1,2 cepdl1,2* quadruple mutant (Noessen) on 3 mM NO_3_^-^. Under these conditions, root nitrate content was similar to wild-type levels (Ota et al., 2020). In our hands, the shoot nitrate content was reduced by a factor of 5.2, while the root nitrate content was increased by a factor of 1.7 (Supplemental Fig. 9). Albeit these supposedly ecotype-specific differences in the relative distribution of nitrate, both data sets are consistent with the notion that root to shoot allocation is severely impaired in the absence of CEPDs.

### TGA1 and TGA4 are CEPD-inactivated activators of genes related to photosynthesis

Transgenic *35S:TGA1* plants ectopically expressing TGA1 were shown to directly or indirectly activate N dose-induced genes in roots of plants that had experienced an N starvation period (Swift et al., 2020). The genes belonged to enriched GO terms “photosynthesis”, “glucose metabolic process” “hexose metabolic process”, “monosaccharide metabolic process” and “generation of precursor metabolites and energy” which were also enriched within the group of 212 genes that were repressed by CEPDs (Supplemental Fig. 5). Expression analysis of four selected genes showed that higher expression levels in *cepd* depend on TGA1/4 (Fig. 4, Supplemental Fig. 10) allowing the assumption that the majority of these genes might be activated by TGA1/4 in the absence of CEPDs. Ectopic expression of TGA1/4 might have out-competed repressive CEPDs at promoters that are activated by TGA1/4.

Enhanced fresh weight of the *35S:TGA1* plants and reduced root fresh weight of *tga1 tga4* might be due to enhanced or compromised expression of at least a subset of these genes. However, this set of genes should be hyper-induced in *cepd*, but the *cepd* mutant has - like *tga1 tga4* - smaller roots. It might well be that processes related to reduced shoot fresh weight and reduced nitrate content in the shoot interfere with the phenotypic consequences of increased expression of TGA1/4-activated growth-promoting genes in *cepd*.

### The CEPD-TGA1/4 regulatory module is not required for nitrate acquisition and shoot growth

As discussed above, the *cepd tga1 tga4* hexuple mutant has the same phenotype as the wild-type when it comes to shoot and root nitrate levels, shoot fresh weight (Figs. 1 and 6) and LN-induced expression of *NRT2.2* and *NRT2.4* upon transfer of plants from FN to LN medium (Fig. 3). This raises the question on the function of the CEPD-TGA1/4 regulatory module. The *cepd* shoot growth phenotype, which is most likely due to impaired translocation of N from the root to the shoot, seems to be independent of the N supply as it became visible when plants were grown on 1 and 10 mM NO_3_^-^ and on fertilized soil. Under all these conditions, *tga1 tga4* did not have an altered shoot fresh weight phenotype, indicating that basal levels of CEPDs are already sufficient to fully repress TGA1/4 activity. Modulation of TGA1/4 activity can thus only be achieved under conditions where even basal CEPD activities are lowered. Comparison of shoot growth of wild-type and *cepd tga1 tga4* under different conditions might unravel the relevance of the CEPD-TGA1/4 regulatory module. Since reduction of the endogenous cytokinin pool reduces transcription of *CEPDs* (Ota et al., 2020; Taleski et al., 2023), environmental or developmental conditions leading to decreased cytokinin levels might lead to reduced shoot growth through increased TGA1/4 activity.

### Regulation of TGA1/4 activity by CEPDs does not involve redox modifications of reactive cysteines

At least class I glutaredoxins can regulate the redox state of thiols in target proteins. Due to their sequence similarity with class I glutaredoxins, CC-type glutaredoxin-like proteins have been suggested to control the redox state of reactive cysteines of their TGA targets. In maize, biochemical evidence suggests that the CC-type glutaredoxin-like proteins MSCA1/ZmGRXC2/ZmGRXC5 negatively control the *in vivo* DNA binding activity of TGA factor FEA4 through keeping Cys321 in the reduced state thus interfering with formation of a dimer-promoting intermolecular disulfide bridge (Yang et al., 2021). However, the *in vivo* importance of Cys321 in FEA4 has not been shown yet. The Arabidopsis TGA factor PAN encodes Cys340 in the corresponding position. Cys340 is important for PAN function *in vivo* (Li et al., 2009), but does not influence the DNA binding activity of the recombinant protein (Gutsche et al., 2017). This is different from FEA4Cys321^Ser^, which fails to bind to DNA. In *Marchantia polymorpha* TGA, Cys231 is at a position equivalent to FEA4Cys321 and PANCys340. The MpTGACys231^Ser^ variant shows enhanced DNA binding activity and is less redox sensitive (Gutsche et al., 2017).

In TGA1 and TGA4, Cys260 corresponds to the cysteines discussed above. Diamide treatment of the recombinant protein led to the formation of an intramolecular disulfide bridge between Cys260 and Cys266, which did not directly affect the DNA binding activity (Despres et al., 2003). Mutating these cysteines had no effect on *in vivo* TGA1 activity with respect to its activating function in the context of hyponasty and defense responses (Li et al., 2019; Budimir et al., 2021) nor with respect to its regulatory function within the N signaling network (Fig. 7). It appears that already the function of the conserved cysteine in different TGA factors varies suggesting that the mechanism of action of CC-type glutaredoxin-like proteins might also vary. Still, the strict conservation of the CCxC/S motif implies a common function of all CC-type glutaredoxin-like proteins which remains to be elucidated.

### CEPDs only slightly influence the activity of TGA1 and TGA4 in the shoot

*CEPDs* are predominantly expressed in the phloem cells of the shoot (Ohkubo et al., 2017; Ota et al., 2020). *TGA1* and *TGA4* expression has also been shown to be more prominent in the vascular tissue than in the in other tissues of the shoot (Wang et al., 2019), and *TGA1* expression is considerably stronger in roots (Supplemental Fig. 3). Under sufficient N supply, TGA1/4 act as transcriptional activators of *ROXY6*, *ROXY8* and *ROXY9* (Fig. 7; Supplemental Fig. 2). *ROXY9* expression is increased in *cepd* (Supplemental Fig. 8) which might be either due to the lack of the repressive activity of CEPDs on TGA1/4 or to the low nitrate content in this mutant.

In shoots, TGA1/4 contribute to the activation of developmental genes like *ARABIDOPSIS THALIANA HOMEOBOX GENE1* (Wang et al., 2019), which is needed for boundary establishment, auxin-inducible genes that are required for cell expansion during hyponastic growth (Li et al., 2019) and genes involved in the synthesis of defense hormones after pathogen infection (Sun et al., 2018). We provided evidence that CEPDs dampen TGA1/4 activity to allow the lowering of petioles after a period of hyponastic growth (Fig. 8). TGA1/4-mediated activation of expression of defense gene *FMO1* was significantly but not strongly counteracted by CEPDs (Fig. 9). Whether lower efficiency of CEPD action in the shoot is due to an unfavorable ratio of CEPDs over TGA1/4 or to other factors missing in the relevant cell types is not known.

Unexpectedly, mutation of the *CEPD* genes in the *tga1 tga4* mutant partially restored upward movement of leaves during the dark period especially if the preceding light period was under low light conditions (Fig. 8b, Supplemental Fig. 13). This indicates that CEPDs can repress hyponasty-promoting responses other than those driven by TGA1/4. A possible scenario might be, that – in the absence of TGA1/4 – other TGAs are targeted to genes involved in hyponasty and that these are repressed by CEPDs.

## Materials and Methods

### Plant material

All plants are in the *Arabidopsis thaliana* Col-0 ecotype background, which was used as a wild-type control. References for published mutants are: *tga1 tga4* (Shearer et al., 2012) and *cepr1-3* (Chapman et al., 2019). Mutant alleles of *ROXY6*, *ROXY7*, *ROXY8*, *ROXY9* and *ROXY10* and deletion of the gene cluster encoding *ROXY11, ROXY12, ROXY13, ROXY14* and *ROXY15* were constructed using a CRISPR/Cas9 system as described below. To obtain the *cepd* (*roxy6-9*) quadruple mutant, *roxy6* and *roxy9* were created first as single mutants in the Col-0 background. The *roxy9* plants were transformed with the particular CRISPR/Cas9 constructs to generate *roxy7,9* and *roxy8,9* double mutants, respectively. After introducing the *roxy6* mutant allele into the *roxy7,9* plants by crossing, the resulting *roxy6,7,9* triple mutant was further crossed with *roxy8,9* enabling the identification of a *roxy6-9* (*cepd*) quadruple mutant. From the analyzed segregating population, a *roxy6,8,9* triple mutant was additionally isolated. The *cepd tga1 tga4* hexuple mutant was obtained by crossing the *cepd* quadruple mutant with the *tga1 tga4* double mutant. Homozygous *roxy10* and *roxy11-15* plants were crossed to create the *roxy10-15* hexuple mutant. Genotyping was done with primers listed in Supplemental Table 2.

### Plant growth conditions

Surface-sterilized seeds were grown in vertically oriented 10 × 10 x 2 cm square plastic Petri dishes (10 seeds per plate). The basal composition of all media was 3 mM KH_2_PO_4_/K_2_HPO_4_, pH 5.8, 4 mM CaCl_2_, 1 mM MgSO_4_, 2 mM K_2_SO_4_, 3 mM MES, pH 5.8, 0.5% (w/v) sucrose, and microelements (i.e. 40 *μ*M Na_2_FeEDTA, 60 *μ*M H_3_BO_3_, 14 *μ*M MnSO_4_, 1 *μ*M ZnSO_4_, 0.6 *μ*M CuSO_4_, 0.4 *μ*M NiCl_2_, 0.3 *μ*M Na_2_MoO_4_, 20 nM CoCl_2_). For growth on full nutrition (FN), 2 mM KNO_3_, 1 mM NH_4_NO_3_, 1 mM L-glutamine, was added, for growth on low N (LN) 0.1 mM KNO_3_, 50 *μ*m NH_4_NO_3_, and 3 mM KCl was added (Scheible et al., 2004). Growth without any reduced N was done in basal media supplemented with either 1 mM KNO_3_/9 mM KCl, 3 mM KNO_3_/7 mM KCl or 10 mM KNO_3_. Petri dishes were stored at 4°C for 24 to 48 hours with the lid facing downwards. For gene expression analysis in shoots or roots under FN and LN doses, plants were first grown for 7 days on FN medium. Subsequently, plants were either transferred to FN or to LN medium and incubated for two days before harvest. For gene expression analysis and determination of nitrate content and shoot and root fresh weight, plants were grown for 21 days. Plants were cultivated at 22°C with continuous light at a photon flux of 70 to 80 µmolꞏm^−2^ꞏs^−1^. For measurements of rosette fresh weight, hyponastic growth analysis and pathogen infections, plants were grown for four weeks on fertilized soil (Ulrich et al., 2021) under different photoperiods as indicated.

### Plant treatments

For endpoint measurements of hyponastic growth in response to low light (LL; 15-20 µmol photons m^2^ s^-1^), we followed the protocols as described before (Li et al., 2019). Briefly, three sets of 8 plants were subjected to the following light regimes. The first set was kept at normal light (12 NL/12 D; 100-120 µmol photons m^2^ s^-1^) throughout the experiment, the second set was subjected to LL conditions, under which they remained for 10.5 hours until the onset of the dark period and for another 4 h after the end of the dark period. The third set was subjected for 10.5 h to LL during the first photoperiod, but was transferred back to NL for 4 h during the second photoperiod. Leaf angle measurements and harvesting of petioles was done at the same time for all plants which is at 4 h after the onset of the second photoperiod. Plants were photographed and the leaf angle of leaf #8 relative to a horizontal line was measured using the angle measurement tool of ImageJ. Hyponastic growth analyses over time were performed with 4-week-old plants grown in NL (9 NL/15 D; 100-120 µmol photons m^2^ s^-1^). Plants were moved at two hours after the dark period (ZT2) to experimental conditions of either NL or LL (approximately 25 µmol photons m^2^ s^-1^). Image acquisition every minute and subsequent analyses were performed as described (Oskam et al., 2023). For analysis of *FMO1* expression upon pathogen infection (Nair et al., 2021), *Pseudomonas syringae* pv. *maculicola* (*Psm*) ES4326 was grown overnight at 28°C in Kings B media supplemented with 50 mg/L rifampicin. Bacteria were washed twice with 10 mM MgCl_2_ and diluted to a final O.D. of 0.005. Three leaves of 4.5-week-old-plants were infiltrated with either 10 mM MgCl_2_ (mock) or *Psm*. Two days after the primary infiltration, three upper leaves were inoculated with *Psm* (O.D. 0.005). Samples were collected eight hours post inoculation (hpi) for gene expression analysis.

### Genome editing

The *roxy* mutants were generated via the clustered regularly interspaced short palindromic repeats (CRISPR)/CRISPR-associated protein 9 (CRISPR/Cas9) genome editing system. The sgRNA targeting sequences (Supplemental Table 3) were cloned into the vector pBCsGFPEE (Nair et al., 2021) for *Agrobacterium*-mediated transformation of *Arabidopsis* plants. If the sgRNA starts with a G, no additional guanine nucleotide serving as RNA polymerase III transcription initiation site was added to the sequences. Homozygous mutants without the CRISPR/Cas9 construct to ensure genetic stability were identified as described in Nair et al., 2021.

The deletions at the *ROXY10* and *ROXY11-15* loci were obtained with T-DNA constructs encoding two sgRNA expression cassettes (Xing et al., 2014), respectively. The particular target sequences were integrated into a PCR product by primers flanking an sgRNA scaffold, the *Arabidopsis thaliana U6-26* snRNA terminator and the *U6-29* promoter sequence as shown in Supplemental Fig. 14. The PCR products were cut with *Bpi*I and *Bsa*I, respectively and cloned into the pBCsGFPEE vector (Nair et al., 2021) cut with *Bsa*I. Since the target sequence 1 is present in both, the *ROXY11* as well as the *ROXY15* locus, sequencing results indicated that the CRISPR/Cas9 editing effect was only mediated by target 1 without modifications at the *ROXY11* specific target 2 site (Supplemental Fig. 1).

### Western blot analysis, RNA extraction, qRT-PCR and RNAseq analysis

Protein extracts and Western blot analysis for detection of TGA1 was done exactly as described (Budimir et al., 2021). RNA extraction, cDNA preparation and qRT-PCR analyses were performed as described previously (Fode et al., 2008). Calculations were done according to the 2^−ΔCT^ method (Livak and Schmittgen, 2001) using the *UBQ5* transcript as a reference (Kesarwani et al., 2007). Primers used to analyze transcript abundance are listed in Supplemental Table 2. RNAse and bioinformatic analysis was done as described (Ulrich et al., 2021) with one exception: reads aligned to the feature type “CDS” instead of “exon” were quantified. Enrichment analysis of TGA factor binding motifs within the 1,000-bp upstream regions was performed as described previously (Berendzen et al., 2012; Zander et al., 2014) using the Cluster Analysis Real Randomization algorithm incorporated into Motif Mapper version 5.2.4.0. Gene Ontology (GO) enrichment analysis was performed using the Gene Onthology Consortium web site ((Ashburner et al., 2000); The Gene Ontology Consortium, 2021) and the PANTHER Classification system version 17.0 (Mi et al., 2013). Differentially expressed genes were sorted according to their biological process into GO groups.

### Determination of nitrate content

Nitrate content determination was based on a spectroscopic method as described (Hachiya and Okamoto, 2017). Briefly, nitrate was extracted from 10 to 50 mg material that was frozen in liquid N_2_. After adding 10 volumes of water, the samples were incubated for 20 min in a 100°C water bath. Cooled tubes were centrifuged for 10 minutes at 20,400 x *g* and 10 µl of the supernatants were incubated with 80 µl of 0.05% (w/v) salicylic acid in sulphuric acid for 20 min at room temperature. 1 ml of 8% NaOH in water was added and the samples were vortexed until the contents became clear. Absorbance was measured at 410 nm along with a calibration curve.

### Statistical analysis

GraphPad Prism 10 (GraphPad Software, Inc., San Diego, CA) was used to conduct statistical analysis.

### Accession numbers

Sequence data from this article can be found in the Arabidopsis Genome Initiative or GenBank/EMBL databases under the following accession numbers:

CEPH (At4g32950); CLE3 (At1g06225); FMO1 (At1g19250); NRT1.1 (At1g12110); NRT1.5 (At1g32450); NRT2.1 (At1g08090); NRT2.2 (At1g08100); NRT2.4 (At5g60770); PER10 (At1g49570); PER71 (At5g65120); ROXY6/GRXS11/CEPD1 (AT1g06830); ROXY7/GRXS9/CEPDL2 (At2g30540); ROXY8/GRXC14/CEPDL1 (At3g62960); ROXY9/GRXC13/CEPD2 (At2g47880); ROXY10/GRXS2 (At5g18600); ROXY11/GRXS3 (At4g15700); ROXY12/GRXS5 (At4g15690); ROXY13/GRXS4 (At4g15680); ROXY14/GRXS7 (At4g15670); ROXY15/GRXS8 (At4g15660); SWEET11 (At3g48740); TFL1 (At5g03840); TGA1 (At5g65210); TGA4 (At5g10030); UBQ5 (At3g62250); UMAMIT35 (At1g60050)

## Supplemental Data

**Supplemental Figure 1** CRISPR/Cas9-based genome editing of *ROXY* genes

**Supplemental Figure 2** Transcript levels of *CEPDs* Col-0, *tga1 tga4* and *cepr1-3* grown under full and limiting N supply

**Supplemental Figure 3** Transcript and protein levels of *TGA1* and *TGA4* in Col-0 grown under full and limiting N supply

**Supplemental Figure 4** Transcript levels of CEPD-activated genes in roots of Col-0 and *cepr1-3* grown under full and limiting N supply

**Supplemental Figure 5** GO term analysis of 212 genes that are higher expressed in *cepd*

**Supplemental Figure 6** Transcript levels of CEPD-repressed genes in roots of Col-0 and *cepr1-3* grown under full and limiting N supply

**Supplemental Figure 7** *UMAMIT35* transcript levels of in roots of Col-0 and *roxy10-15* grown under full nitrogen supply

**Supplemental Figure 8** Transcript levels of N starvation-induced genes in of Col-0, *tga1 tga4, cepd* and *cepd tga1 tga4* grown on 1 or 10 mM nitrate

**Supplemental Figure 9** Nitrate content in roots and shoots of Col-0, *tga1 tga4, cepd* and *cepd tga1 tga4* grown on 3 mM nitrate

**Supplemental Figure 10** Comparison of *NRT2.2*, *ROXY8* and *ROXY9* transcript levels in plants grown on FN or 10 mM NO_3_^-^

**Supplemental Figure 11** Transcript levels of TGA1/4-activated gene *PER71* in roots of Col-0, *tga1 tga4*, *cepd* and *cepd tga1 tga4* grown under full and limiting N supply

**Supplemental Figure 12** Images of hyponasty of wild-type (Col-0), *tga1 tga4, cepd* and *cepd tga1 tga4* plants

**Supplemental Figure 13** Kinetics of hyponastic growth of *tga1 tga4* and *cepd tga1 tga4* plants

**Supplemental Figure 14** PCR products used for CRISPR/Cas9-based genome editing of the *ROXY10* and *ROXY11-15* loci

**Supplemental Table 1** List of differentially expressed genes (Col-0 vs. *cepd*)

**Supplemental Table 2** List of primers for genotyping

**Supplemental Table 3** Single guide RNA (sgRNA) targeting sequences

**Supplemental Table 4** Primers for qRT-PCR

## Acknowledgments

We acknowledge funding from the German Research Foundation (GRK 2172-ProTECT). We thank Anna Hermann, Katharina Dworak and Ronald Scholz (all UNI-GOE) for excellent technical assistance, Hannah Knerich and Lena Zanoni (both UNI-GOE) for help with gene expression analysis and Florian Jung (UNI-GOE) for help when generating the *roxy9* mutant. We also thank the Next Generation Sequencing-Integrative Genomics Core Unit (NIG), Institute of Human Genetics, at the University Medical Centre Göttingen (UMG), Germany, for performing the RNAseq analysis.

## Author contributions

C.T. and C.G. conceptualized the study. C.T. generated the CRISPR mutants and analyzed transcriptome data. A.M.P. performed the experiments involving gene expression analysis upon transfer of plants from LN to FN media and pathogen attack, supervised the generation of the *cepd tga1 tga4* mutant and did end-point determinations of hyponasty. B.H. performed the analysis of plants grown on media containing various concentrations of nitrate. J.B. helped with the generation of the CRISPR mutants and performed Western blot analysis. L.O. performed time-resolved hyponasty experiments under the supervision of R.P.. C.G. wrote the manuscript. All authors helped in writing and revising the manuscript.

---

**Fig. S1.**
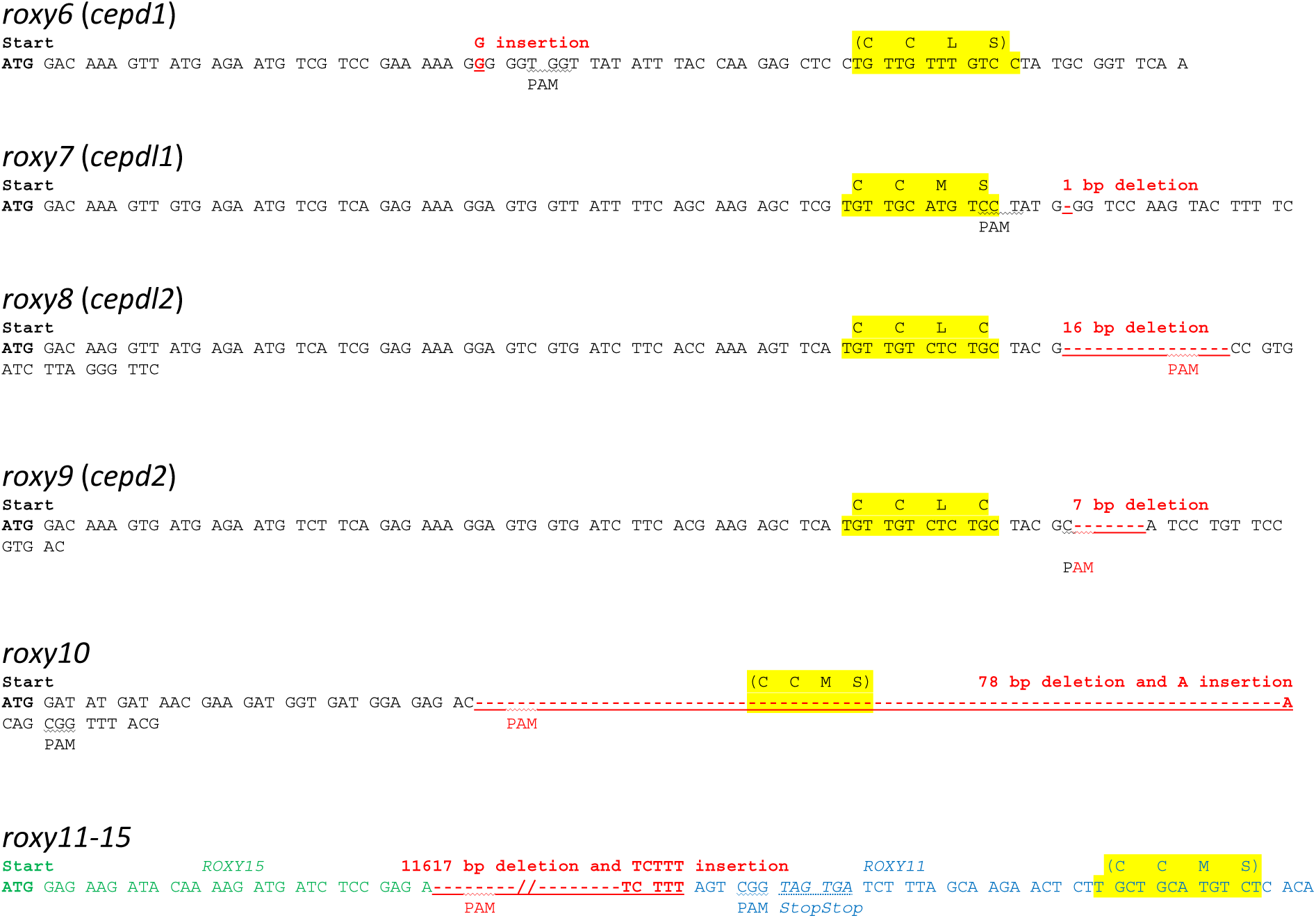
CRISPR/Cas9-based genome editing of *ROXY* genes. The active site-encoding sequence is highlighted in yellow. Mutations leading to frameshifts and deletions are indicated in red. PAM, protospacer adjacent motif.

**Fig. S2.**
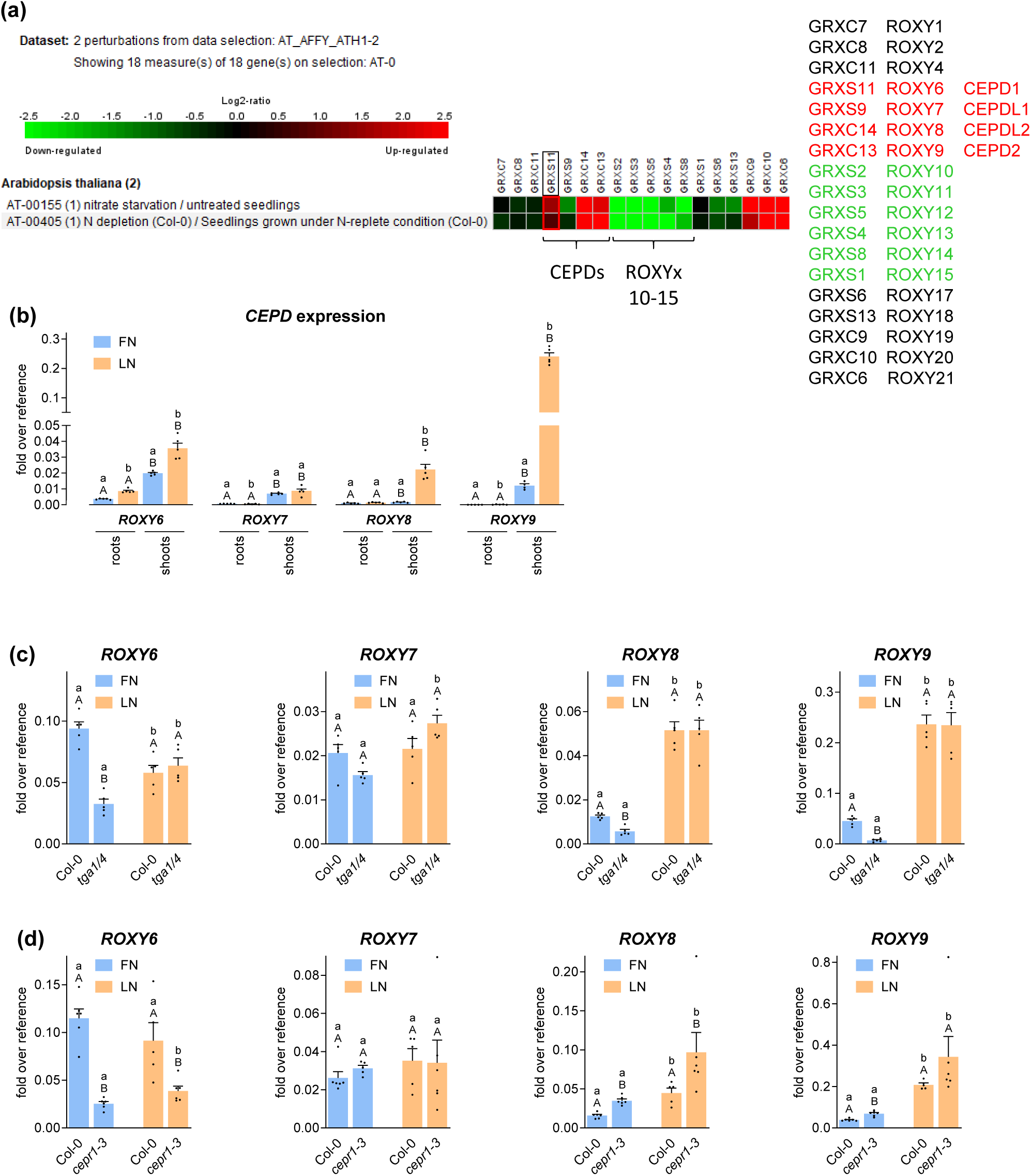
Transcript levels of *CEPD*s in Col-0, *tga1 tga4* and *cepr1-3* grown under full and limiting N supply. **(a)** Expression levels of CC-type glutaredoxins according to www.genevestigator.de. Seven-day-old seedlings grown in liquid full nutrition (FN) medium under constant light were transferred to either FN or low nitrogen (LN) medium. Two days later, seedlings were collected for transcriptome analysis (Scheible et al., 2004). **(b-d)** Seedlings were grown as in **(a),** but on agar-solidified medium. Shoots and roots were collected for RNA isolation, in **(c)** and **(d)**, only shoot RNA was analysed. Expression of the indicated genes was analysed by qRT-PCR, *UBQ5* was used as a reference gene. Mean values of four to five biological replicates are shown, with one replicate originating from one plate with 10 plantlets. Error bars represent the standard error of the mean. In **(b)**, lowercase letters indicate statistically significant differences within the tissue between the conditions, uppercase letters indicate significant differences within treatment between the tissues. In **(c)** and **(d)**, lowercase letters indicate statistically significant differences within the genotype between the treatments, uppercase letters indicate significant differences within treatment between the genotypes. Statistical analyses were performed with the logarithmic values by using two-way ANOVA and Fisher’s Least Significant Difference test (*p* < 0.05).

**Fig. S3.**
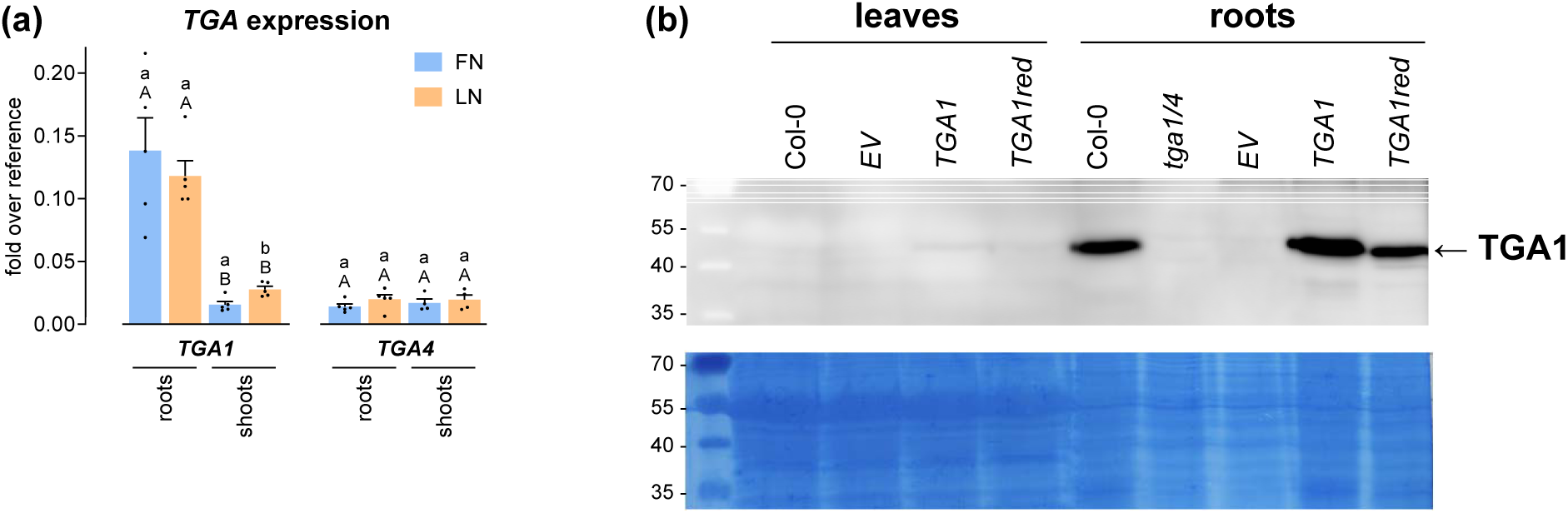
Transcript and protein levels of *TGA1* and *TGA4* in shoots and roots of Col-0 plants. **(a)** 7-day-old seedlings grown on full nitrogen (FN) medium were transferred to either FN or low nitrogen (LN) plates. Two days later, shoots and roots were collected for RNA isolation. Expression of the indicated genes was analyzed by qRT-PCR, *UBQ5* was used as a reference gene. Mean values of four to five biological replicates are shown with one replicate originating from one plate with 10 plantlets. Error bars represent the standard error of the mean. Lowercase letters indicate statistically significant differences within the tissue between the conditions, uppercase letters indicate significant differences within treatment between the tissues. Statistical analyses were performed with logarithmic values by using two-way ANOVA and Fisher’s Least Significant Difference test (*p* < 0.05). **(b)** Western blot analysis of protein extracts obtained from leaves and roots of four-week-old Col-0 and *tga1 tga4* plants complemented either with a control vector *(EV),* a wildtype *TGA1* genomic construct (*TGA1*) or a mutated *TGA1* genomic construct carrying mutations in four critical cysteine residues (*TGA1red*). TGA1 protein levels were detected using an anti-TGA1 antibody. Plants were grown on fertilized soil. Coomassie blue staining served as a loading control.

**Fig. S4.**
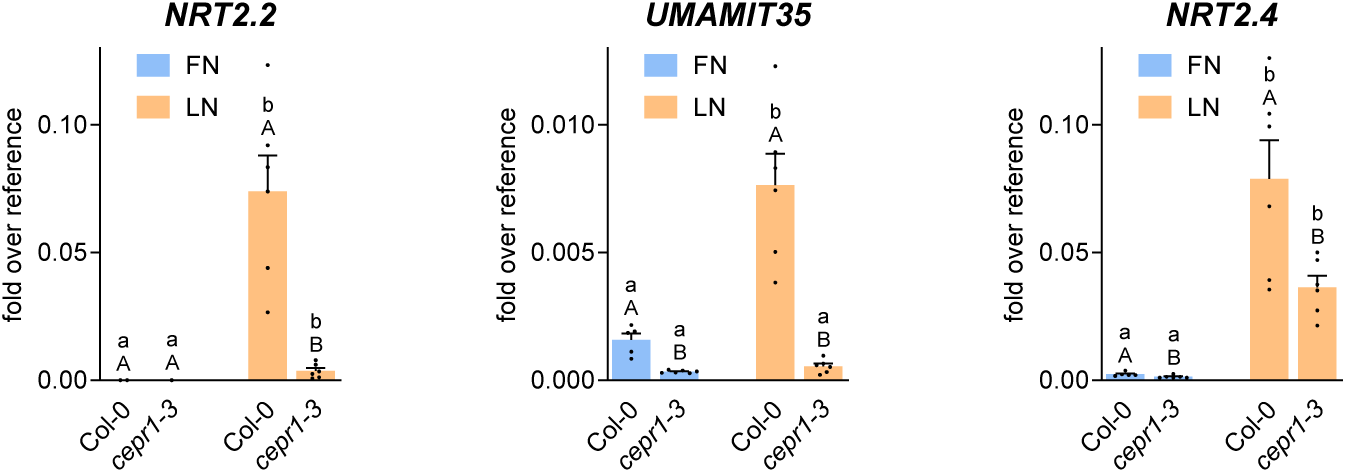
Transcript levels of CEPD-activated genes in roots of Col-0 and *cepr1-3* grown under full and limiting N supply. 7-day-old seedlings grown on full nitrogen (FN) medium were transferred to either FN or low nitrogen (LN) plates. Two days later, roots were collected for RNA isolation. Expression of the indicated genes was analyzed by qRT-PCR, *UBQ5* was used as a reference gene. Mean values of four to five biological replicates are shown with one replicate originating from one plate with 10 plantlets. Error bars represent the standard error of the mean. Lowercase letters indicate statistically significant differences within the genotype between the treatments, uppercase letters indicate significant differences within treatment between the genotypes. Statistical analyses were performed with logarithmic values by using two-way ANOVA and Fisher’s Least Significant Difference test (*p* < 0.05).

**Fig. S5.**
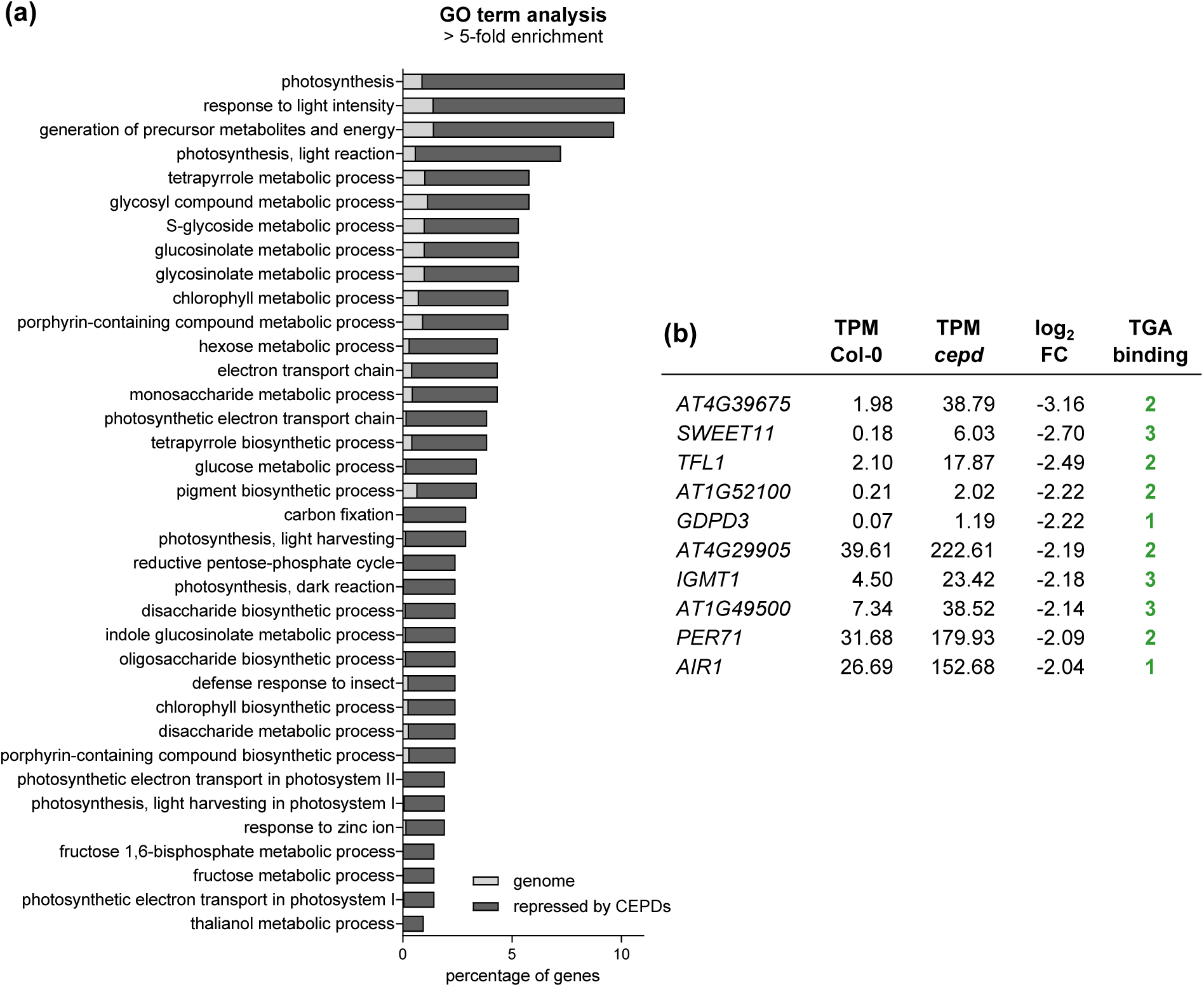
GO term analysis of 212 genes that are higher expressed in *cepd*. **(a)** Gene Ontology (GO) term analysis (biological processes) of 212 differentially expressed genes (log_2_ FC < −1, *p* adj < 0.05) in roots of 9-day-old *cepd* seedlings compared to Col-0 after two days of cultivation on LN (low nitrogen) medium. Bars represent the percentage of genes found per GO term in the group of 212 genes (dark grey) and the percentage of genes representing the respective GO term found within the Arabidopsis genome (light grey). GO terms with > 5-fold enrichment against the genome are shown. Statistical analysis was performed using Fisher’s Exact test and False Discovery Rate (FDR) < 0.05. **(b)** List of the ten most highly differentially expressed genes. The number of TGA binding sites (TGACG and TACGTA) was counted in the region 2 kb upstream of the transcriptional start site. TPM: transcripts per million.

**Fig. S6.**
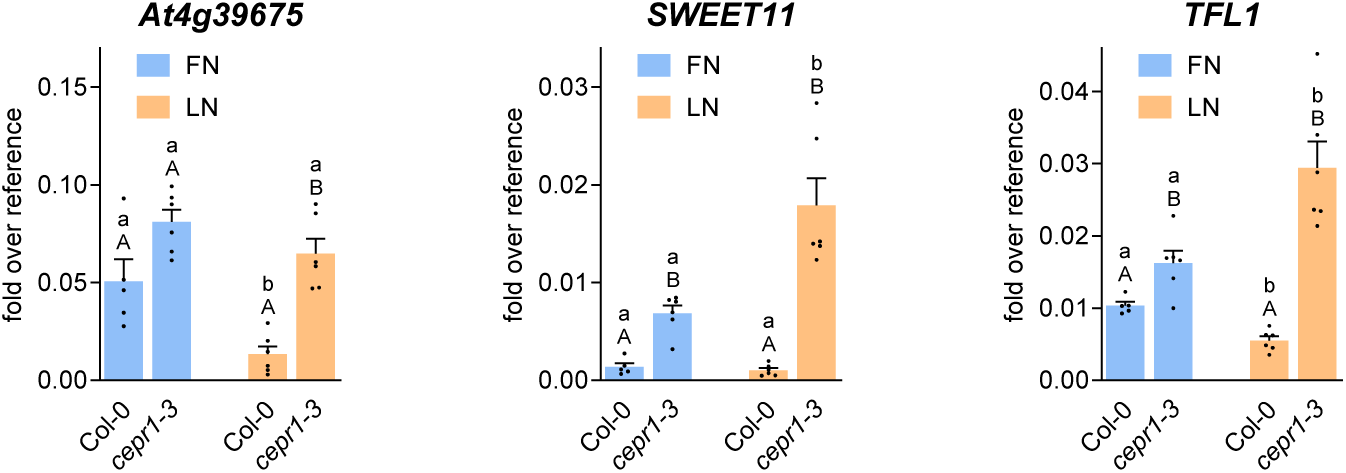
Transcript levels of CEPD-repressed genes in roots of Col-0 and *cepr1-3* grown under full and limiting N supply. 7-day-old seedlings grown on full nitrogen (FN) medium were transferred to either FN or low nitrogen (LN) plates. Two days later, roots were collected for RNA isolation. Expression of the indicated genes was analyzed by qRT-PCR, *UBQ5* was used as a reference gene. Mean values of four to five biological replicates are shown with one replicate originating from one plate with 10 plantlets. Error bars represent the standard error of the mean. Lowercase letters indicate statistically significant differences within the genotype between the treatments, uppercase letters indicate significant differences within treatment between the genotypes. Statistical analyses were performed with logarithmic values by using two-way ANOVA and Fisher’s Least Significant Difference test (*p* < 0.05).

**Fig. S7.**
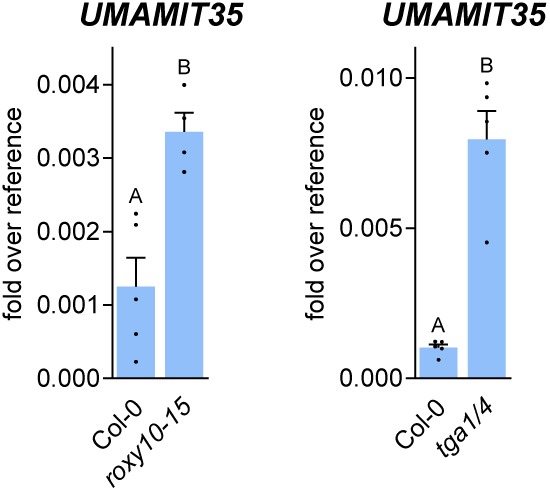
*UMAMIT35* transcript levels of in roots of Col-0 and *roxy10-15* plants grown under full nitrogen supply. 7-day-old seedlings grown on full nitrogen (FN) medium were transferred to FN plates. Two days later, roots were collected for RNA isolation. Expression of the indicated genes was analyzed by qRT-PCR, *UBQ5* was used as a reference gene. Mean values of four to five biological replicates are shown with one replicate originating from one plate with 10 plantlets. Error bars represent the standard error of the mean. Letters indicate significant differences between the genotypes. Statistical analyses were performed with logarithmic values by unpaired t-test (*p* < 0.05). Data from the *tga1 tga4* mutant are the same as in Fig. 2.

**Fig. S8.**
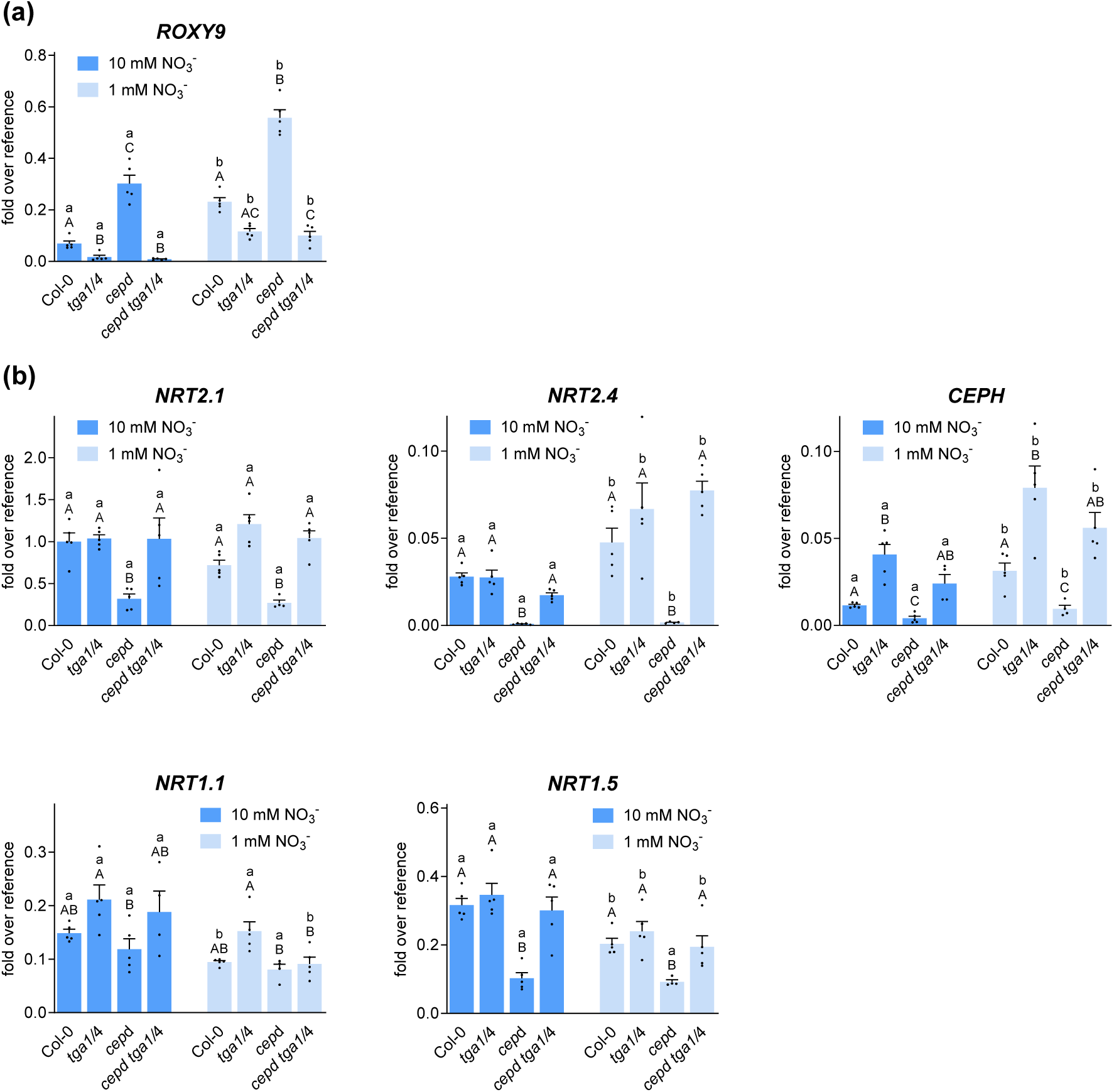
Transcript levels of N starvation-induced genes in Col-0, *tga1 tga4, cepd* and *cepd tga1 tga4* grown on 1 or 10 mM nitrate. Wild-type (Col-0), *tga1 tga4, cepd* and *cepd tga1 tga4* plants were grown for 21 days on medium containing 1 or 10 mM NO_3_^-^ under constant light (70 µmol photons s^-1^ m^- 2^). Transcript levels of *ROXY9* and transcript levels of *NRT2.1*, *NRT2.4*, *CEPH*, *NRT1.1* and *NRT1.5* were determined in shoots and roots, respectively. Mean values of five biological replicates are shown, with one replicate originating from one plate with 10 plantlets. Error bars represent the standard error of the mean. Lowercase letters indicate statistically significant differences within the genotype between the treatments, uppercase letters indicate significant differences within treatment between the genotypes. Statistical analyses were performed logarithmic values by using two-way ANOVA and Bonferroni’s post-test (*p* adj. < 0.05).

**Fig. S9.**
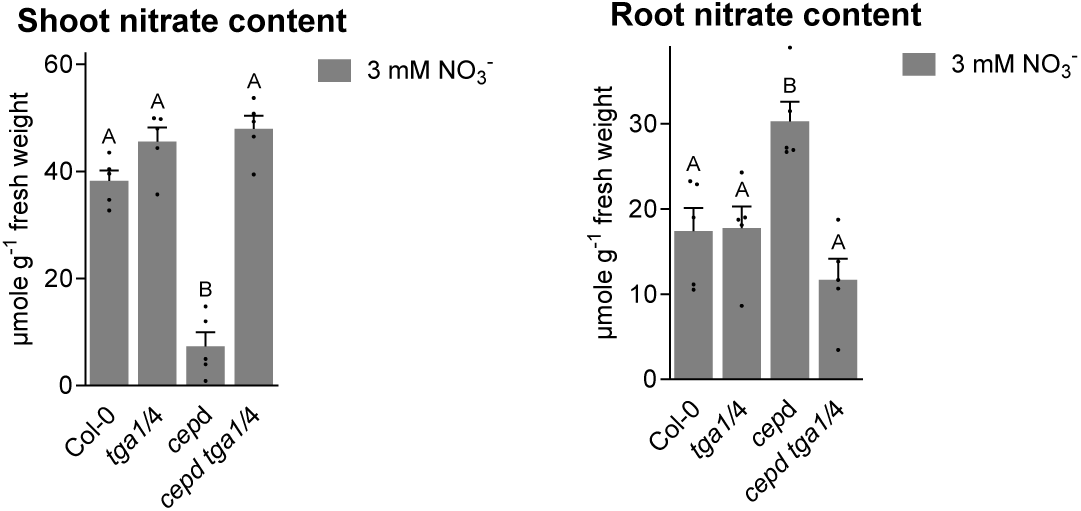
Nitrate content in roots and shoots of Col-0, *tga1 tga4, cepd* and *cepd tga1 tga4* grown on 3 mM nitrate. Wild-type (Col-0), *tga1 tga4, cepd* and *cepd tga1 tga4* plants were grown on medium containing 3 mM NO_3_^-^ under constant light (70 µmol photons s^-1^ m^-2^). After 21 days, shoots and roots were collected separately for determination of the nitrate content. Mean values of five biological replicates are shown, with one replicate originating from one plate with 10 plantlets. Error bars represent the standard error of the mean. Letters indicate significant differences between the genotypes. Statistical analyses were performed with logarithmic values by using one-way ANOVA and Tukey’s post-test (*p* adj. < 0.05).

**Fig. S10.**
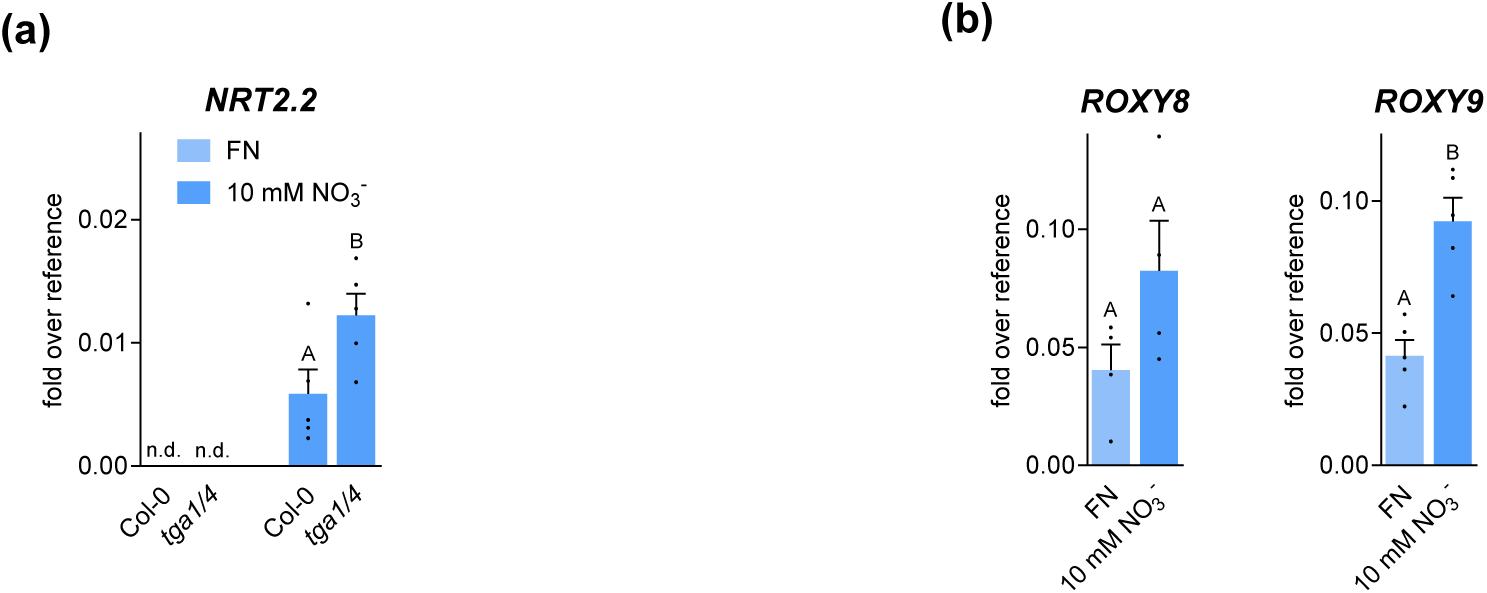
Comparison of *NRT2.2*, *ROXY8* and *ROXY9* transcript levels in plants grown on FN or 10 mM NO_3_^-^. Wild-type (Col-0) plants were grown on media containing 3 mM NO_3_^-^, 1 mM NH_4_ ^+^ and 1 mM glutamine (FN) or 10 mM NO_3_^-^ as the sole N source under constant light (70 µmol photons s^-1^ m^- 2^). After 9 days, shoots and roots were collected separately for analyses of **(a)** *NRT2.2* (roots) **(b)** *ROXY8* and *ROXY9* (shoots) transcript levels. Mean values of (four to) five biological replicates are shown, with one replicate originating from one plate with 10 plantlets. Error bars represent the standard error of the mean. Letters indicate significant differences between the growth conditions. Statistical analysis was performed with logarithmic values by unpaired t-test (p < 0.05).

**Fig. S11.**
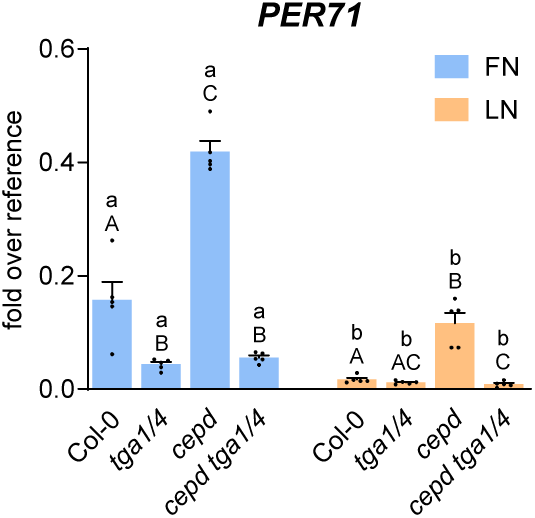
Transcript levels of TGA1/4-activated gene *PER71* in roots of Col-0, *tga1 tga4*, *cepd* and *cepd tga1 tga4* grown under full and limiting N supply. Seven-day-old seedlings grown on full nitrogen (FN) medium were transferred to either FN or low nitrogen (LN) plates. Two days later, roots were collected for RNA isolation. Expression was analyzed by qRT-PCR, *UBQ5* was used as a reference gene. Mean values of four to five biological replicates are shown with one replicate originating from one plate with 10 plantlets. Error bars represent the standard error of the mean. Lowercase letters indicate statistically significant differences within the genotype between the treatments, uppercase letters indicate significant differences within treatment between the genotypes. Statistical analyses were performed with logarithmic values by using two-way ANOVA and Bonferroni’s post-test (*p* adj. < 0.05).

**Fig. S12.**
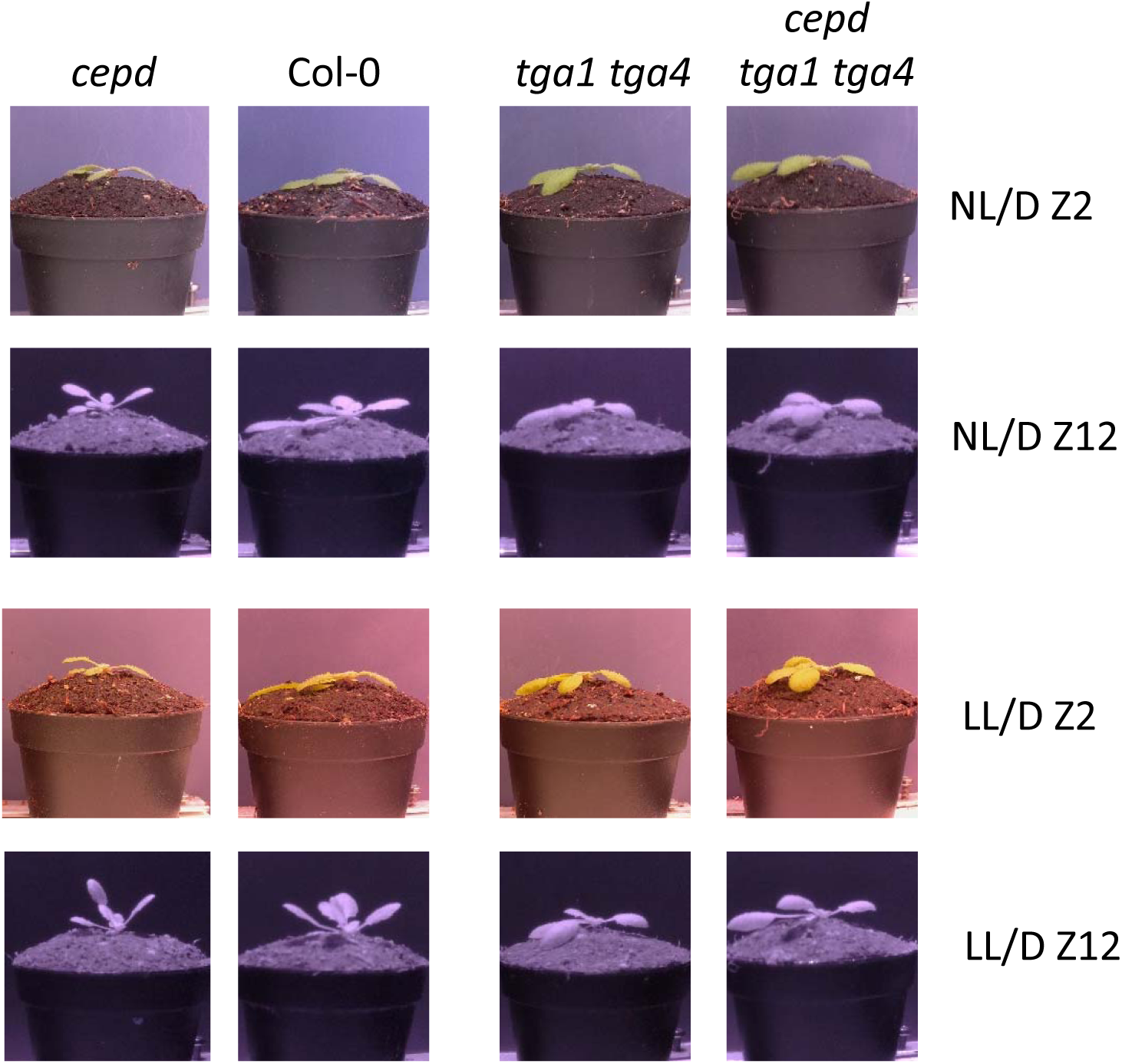
Images of hyponasty of wild-type (Col-0), *tga1 tga4, cepd* and *cepd tga1 tga4* plants. Images belong to the experiments shown in Fig. 8 (Col-0; *cepd*) and Supplemental Figure 13 (*tga1 tga4; cepd tga1 tga4*).

**Fig. S13.**
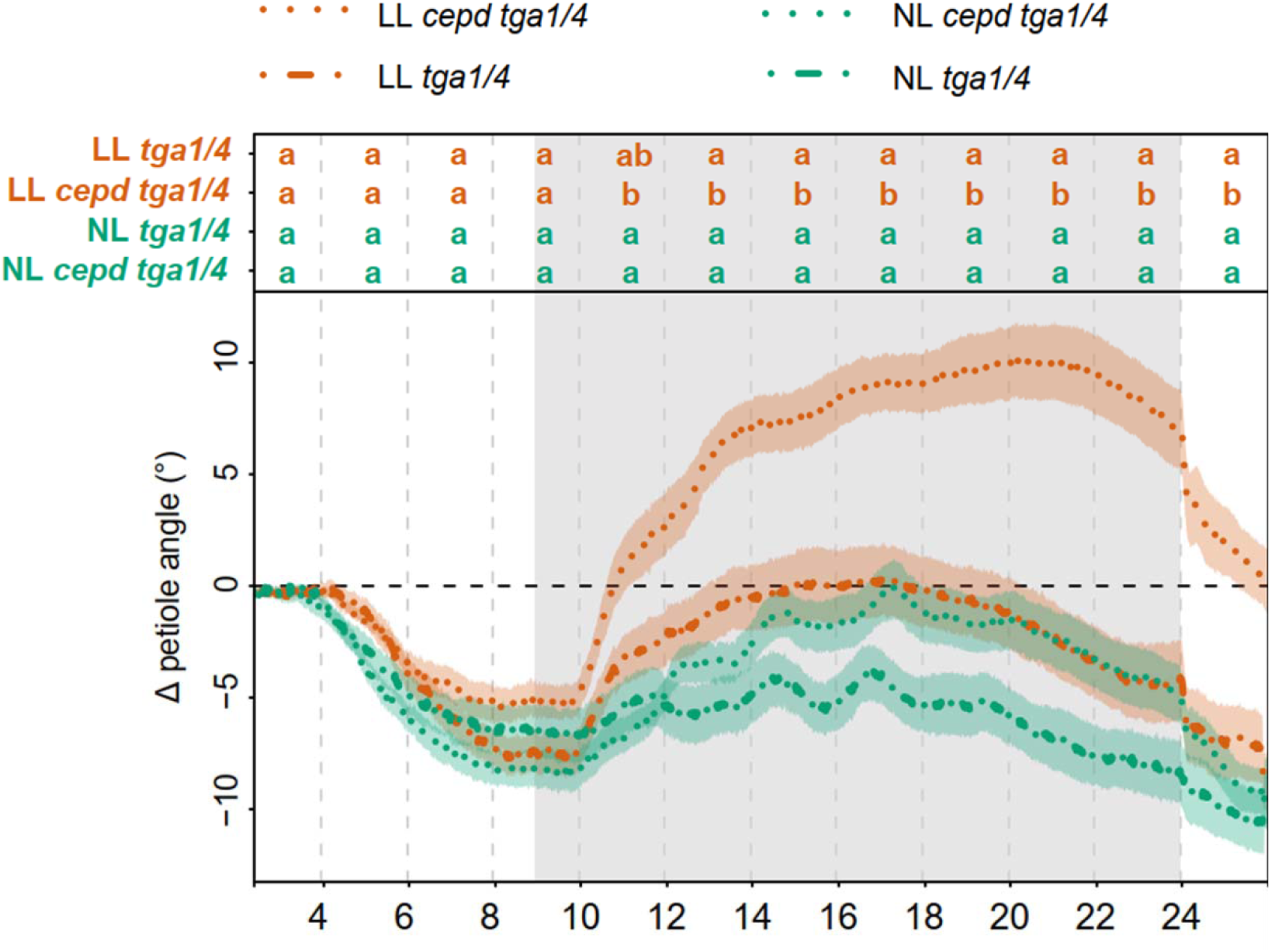
Kinetics of hyponastic growth of *tga1 tga4* and *cepd tga1 tga4* plants. Plants were grown under an NL (100-120 µmol photons m^−2^ s^−1^)/D cycle. Petiole angles of the different genotypes were set to 0 at ZT=2 and the relative angle change was measured every minute for 24 h. At ZT=2, plants were transferred to LL (approximately 25 µmol photons m^−2^ s^−1^). Grey areas indicate the dark period. Letters indicate *p* adj. < 0.05, calculated per every two hours using one-way ANOVA and Tukey post-test n=4).

**Fig. S14.**
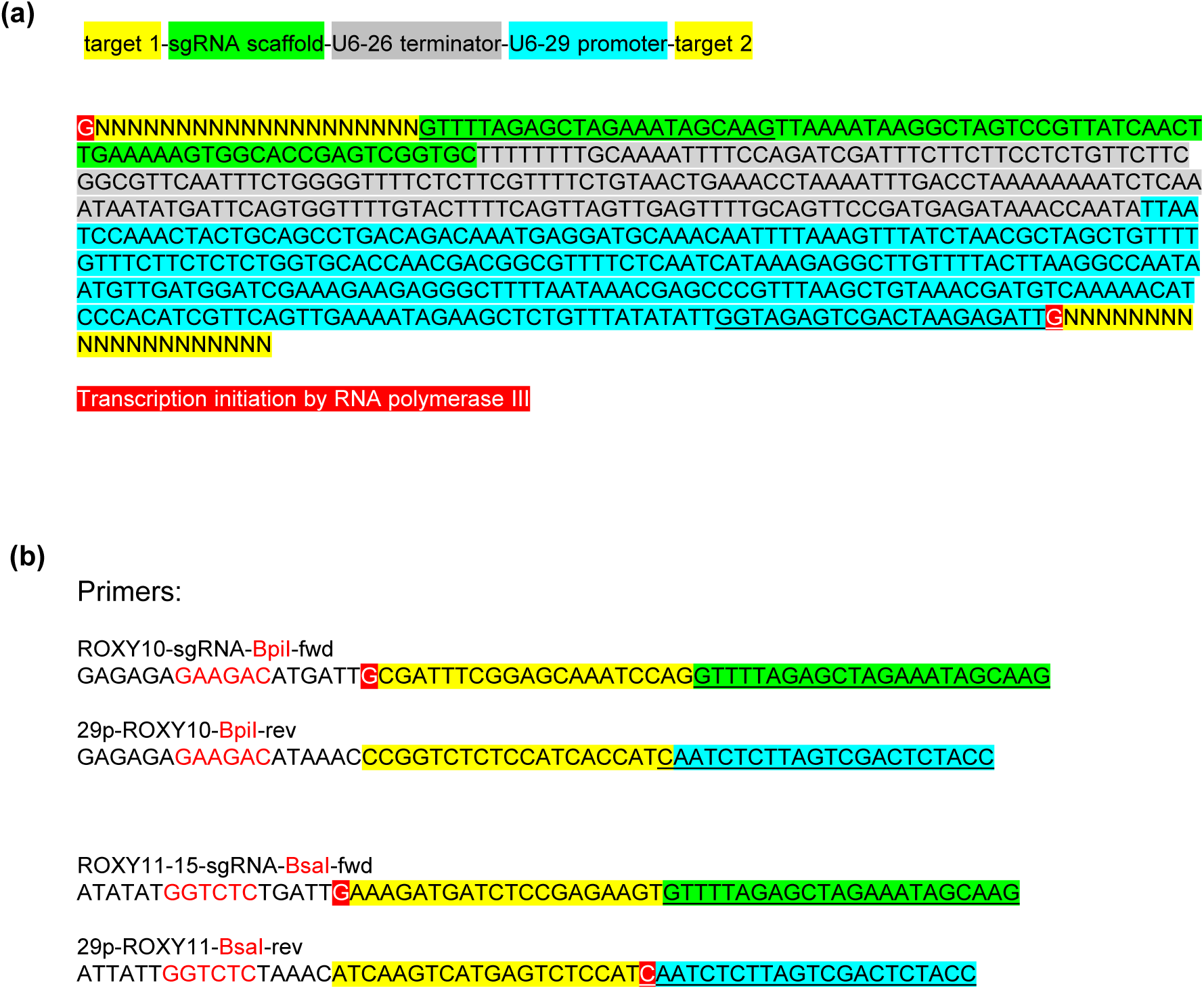
PCR products used for CRISPR/Cas9-based genome editing of the *ROXY10* and *ROXY11-15* loci. **(a)** Scheme and sequence of the PCR product generated to clone two sgRNA expression cassettes (Xing et al., 2014) into the pBCsGFPEE vector (Nair et al., 2021). **(b)** Primers used to introduce the *roxy10* and *roxy11-15* mutations shown in Supplemental Fig. S1, respectively.

**Supplemental Table S2:**
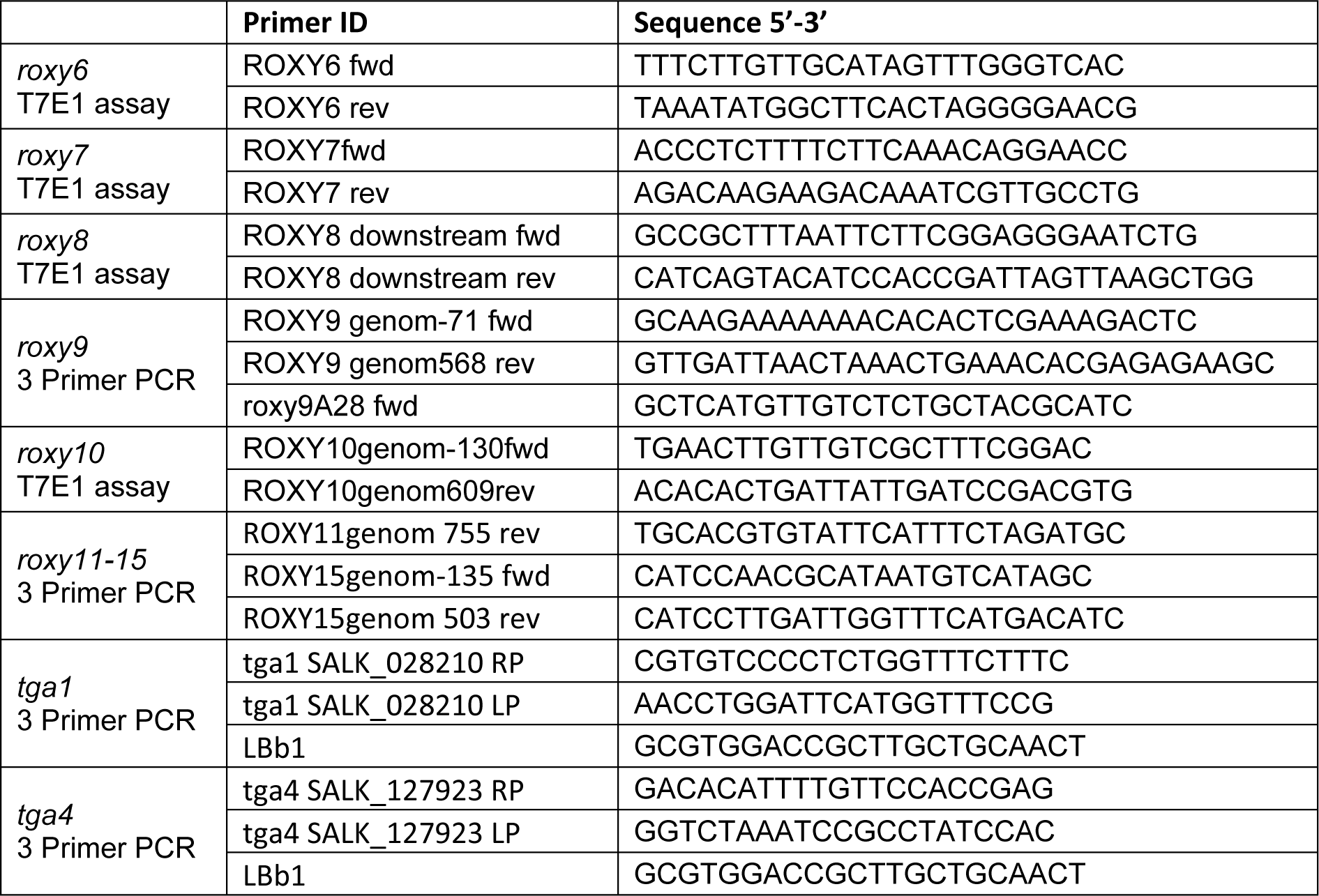
Primers for genotyping.

**Supplemental Table S3:**
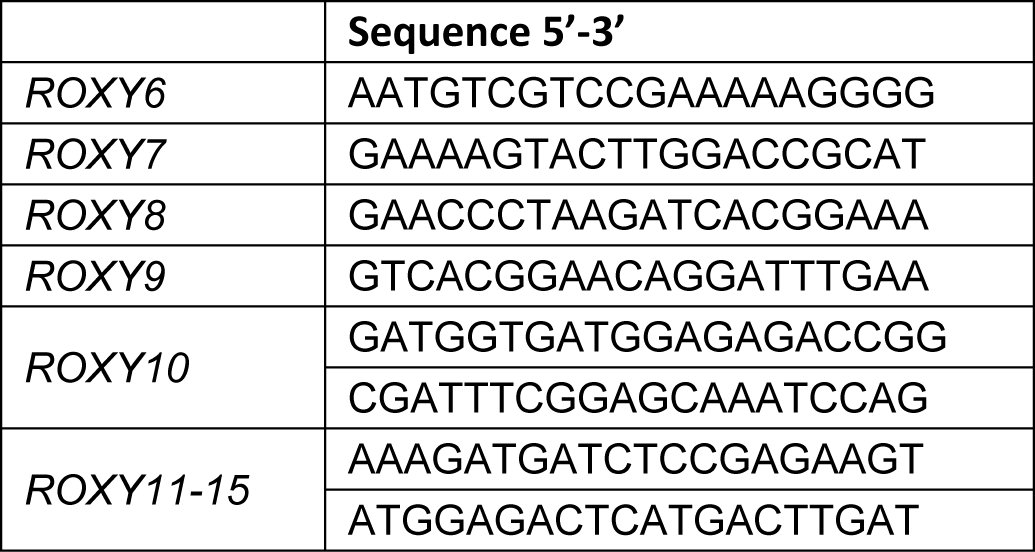
Single guide RNA (sgRNA) targeting sequences.

**Supplemental Table S4:**
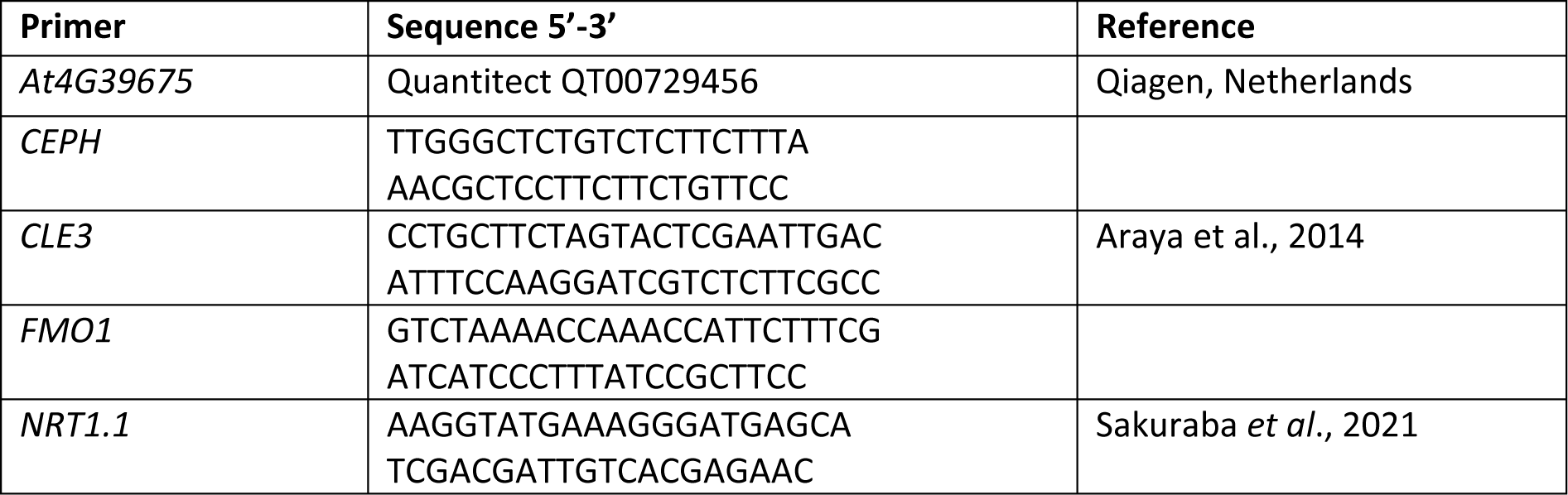
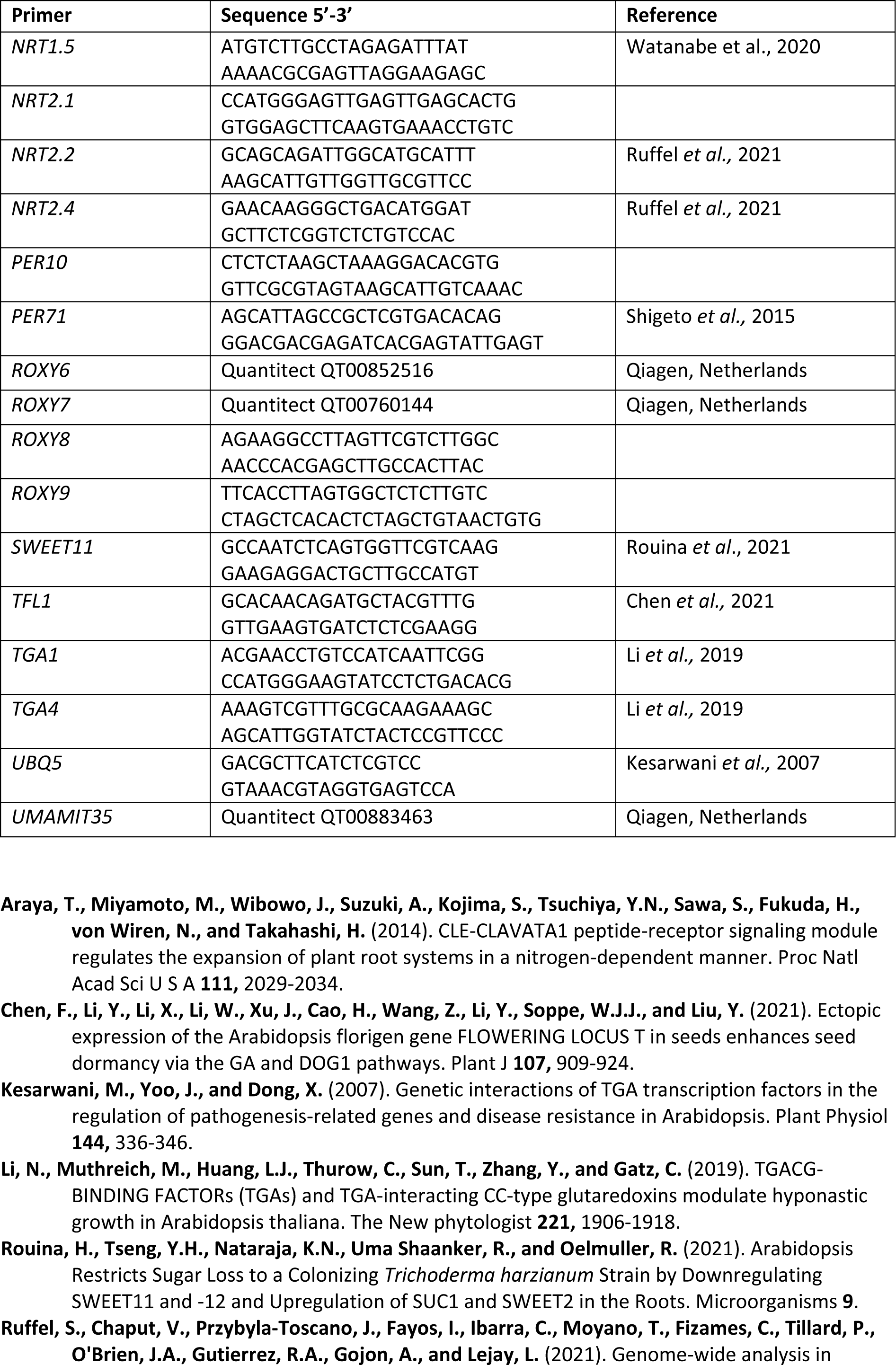
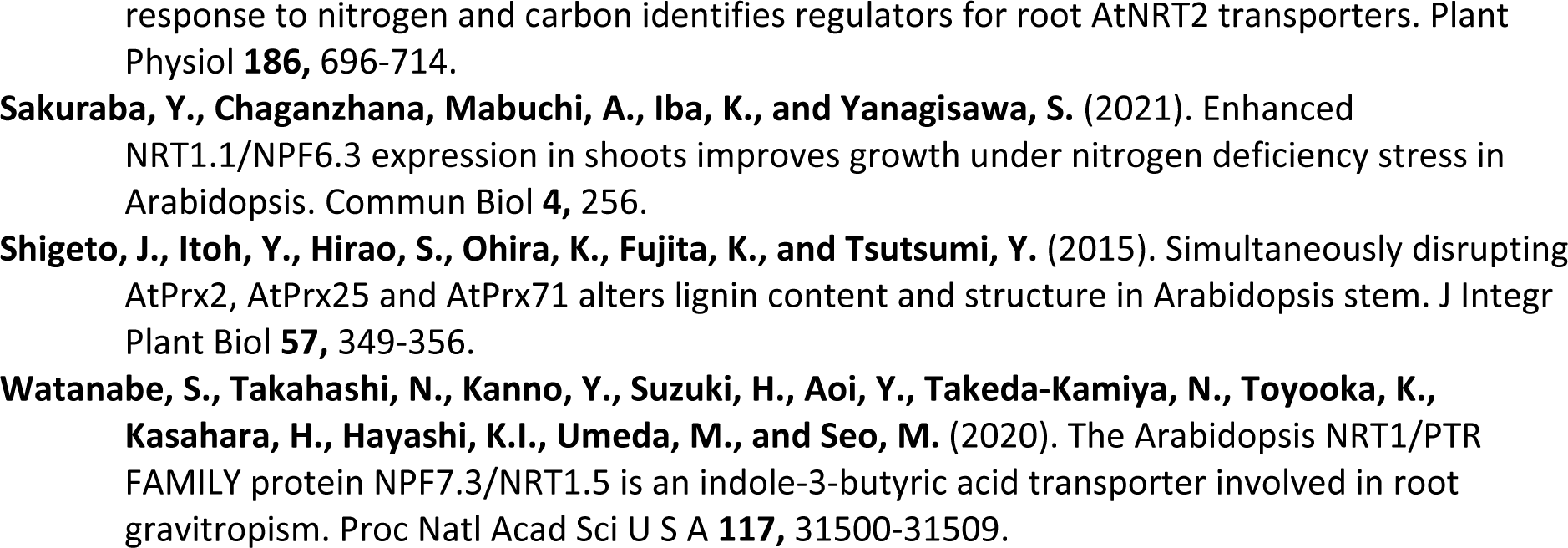
Primers for qRT-PCR.

